# Distinct function of SPL genes in age-related resistance in *Arabidopsis*

**DOI:** 10.1101/2022.12.22.521518

**Authors:** Lanxi Hu, Peng Qi, Alan Peper, Feng Kong, Yao Yao, Li Yang

## Abstract

In plants, age-related resistance (ARR) refers to a gain of disease resistance during shoot or organ maturation. ARR associated with vegetative phase change, a transition from juvenile to adult stage, is a widespread agronomic trait affecting resistance against multiple pathogens. How innate immunity in a plant is differentially regulated during successive stages of shoot maturation is unclear. In this work, we found that *Arabidopsis thaliana* showed ARR against its bacterial pathogen *Pseudomonas syringae pv. tomato* DC3000 during vegetative phase change. The timing of the ARR activation was associated with a temporal drop of miR156 level. A systematic inspection of the loss- and gain-of-function mutants of 11 *SPL* genes revealed that a subset of *SPL* genes, notably *SPL2, SPL10*, and *SPL11*, activated ARR in adult stage. The immune function of SPL10 was independent of its role in morphogenesis. Furthermore, the SPL10 mediated an age-dependent augmentation of the salicylic acid (SA) pathway partially by direct activation of *PAD4*. Disrupting SA biosynthesis or signaling abolished the ARR against *Pto* DC3000. Our work demonstrated that the miR156-SPL10 module in *Arabidopsis* is deployed to operate immune outputs over developmental timing.

**Significance:** Age-associated change of immunity is a widespread phenomenon in animals and plants. How organisms integrate immune maturation into a developmental clock is a fundamental question. Heterochronic microRNAs are key regulators of developmental timing. We found that a conserved heterochronic microRNA (miRNA) in *Arabidopsis*, microRNA156, regulates the timing of age-related resistance associated with a transition from the juvenile to the adult vegetative phase. The coordination between developmental maturation and gain of disease resistance is achieved through miR156-controlled SPL transcription factors with distinct functions. A subset of SPL transcription factors promoted resistance by directly activating key genes in defense signaling. This work bridges the knowledge gap between vegetative development and age-related resistance. Pinpointing mechanisms of the developmental regulation on immunity may pave a way for unlocking the age limit on plant immunity and lay a foundation to applications in the precision agriculture.

## Introduction

Both animals and plants suffer from infectious diseases, particularly at a young age (1-3). The function of their immune systems can be enhanced with the progression of organismal maturation. In many plant species, a gain of disease resistance against certain pathogens during shoot maturation is termed age-related resistance (ARR). Plant ARR can launch robust and wide spectrum resistance against a variety of pathogens, and such trait is often selected in breeding (4).

Age-associated disease resistance is often coupled with successive developmental transitions, such as germination (5) and vegetative phase change (6) and flowering (7, 8). The heterogeneity of host age, maturing stage of infected organs and virulence of causal pathogens suggest that multiple layers of signaling are intertwined between aging and immunity (4, 9). ARR-associated juvenile-to-adult vegetative phase change (hereafter ARR_VPC_) has been observed among economically important vegetables, crops and fruits, such as tomato, rice and grapevine (10). The juvenile and adult phases refer to vegetative development prior to floral induction, and predicable changes of morphological and physiological traits are associated with this transition (11-13). Several factors were speculated to impact ARR_VPC_. Compared to juveniles, adult plants are exposed to environmental conditions that are not optimal for disease development (such as high UV) (3); adult tissues may carry tough physical barriers (e.g., cell wall components, cuticle) (14); and leaves of adult stage are primed by previous exposure to pathogens (3). Such factors complicate the investigation of intrinsic molecular mechanisms governing the onset of ARR_VPC_. Nevertheless, accumulating evidence suggests that intrinsic signaling pathways govern ARR (4, 15).

MicroRNA156s (miR156), a conserved microRNA family (11), regulates the onset of vegetative phase change (11, 16). MiR156 targets genes encoding SPLs (SQUAMOSA PROMOTER BINDING PROTEIN-LIKE) transcription factors, which contains a SQUAMOSA promoter binding protein (SBP) box for nuclear import and DNA binding (17-19). In *Arabidopsis*, leaves generated in the juvenile shoot, e.g., juvenile leaves (usually leaves 1-7 under a short-day condition) accumulate high level of miR156 (20, 21). Throughout the expansion of juvenile leaves, they maintain the morphological (e.g. no abaxial trichome) and molecular identities (e.g., high miR156 level) of the juvenile fate. A temporal decline of miR156 level, followed by the high expression of *SPL*s, initiates the vegetative phase change (19-22). SPLs have overlapping yet distinct functions to promote adult traits such as adult leaf morphogenesis, floral induction, and reduced rooting (23-25). A total of 11 *SPL* genes encoded in *Arabidopsis* Columbia-0 (Col-0) ecotype are suppressed by miR156 via mRNA cleavage and/or translational repression (19, 26). Recent studies showed that the miR156-SPL pathway involved in plant immunity. Disrupting miR156 function in *Arabidopsis* by mutating SQUINT (SQN), an *Arabidopsis* orthologue of cyclophilin 40, compromised jasmonic acid signaling and disease resistance against necrotrophic pathogen *Botrytis cinerea* (27). Furthermore, overexpressing miR156-targeted *SPL9* in juvenile plants enhanced accumulation of reactive oxygen species and induced salicylic acid (SA) signaling, leading to enhanced resistance of *Arabidopsis* against bacterial pathogen *Pseudomonas syringae* (28). Yet, a systematic dissection of the link between ARR and miR156-SPL signaling pathway is still lacking.

Here, we systemically analyzed the miR156-SPLs module in ARR_VPC_. We demonstrated that the ARR to *Pseudomonas syringae pv. tomato* DC3000 (*Pto* DC3000) in *Arabidopsis* is associated with vegetative phase change. Altering the temporal expression of miR156-SPL pathway was sufficient to change the timing of ARR_VPC_ onset. A sub-class of SPL transcription factors (SPL2/10/11) promoted disease resistance in adult stage, and such function was independent of their roles in leaf morphology. Transcriptomic analysis unveiled multiple mechanisms that collectively contribute to ARR_VPC_, including priming, activating adult-specific defense programs, and strengthening juvenile defense after infection. Finally, we found SPL10 strengthened SA signaling in the adult stage by directly enhancing the transcription of PAD4. Our work provides molecular insights into the intrinsic clock that coordinates disease resistance outcomes with developmental timing.

## Results

### The age-related resistance to *Pto* DC3000 is associated with a reduction of miR156 level

To assess the ARR_VPC_, we measured the multiplication of *Pto* DC3000 in juvenile (without abaxial trichomes) and adult (with abaxial trichomes) leaves of *Arabidopsis thaliana* Col-0 ecotype. During the expansion of an individual leaf, defense gene is differentially expressed (SI Appendix, Fig. S1), which is known as ontogenic resistance (5, 29-32). Ontogenic resistance occurs during the maturation of both juvenile and adult leaves (SI Appendix, Fig. S1). Because juvenile leaves are produced in early shoot development, juvenile and adult leaves on a same plant are always at different ontogenic age (SI Appendix, Fig. S1). To avoid the impact of ontogenic resistance, we sampled fully expanded juvenile leaves (leaves 1-4) and adult leaves (leaves 8) from 5-, 6- and 7-week-old plants, respectively. To avoid the influence of flowering-associated ARR (33, 34), plants were grown in a short-day condition and bolting was not observed before 10 weeks after planting. Bacterial multiplication in leaves 8 was lower than that in leaves 1, 2, 3 and 4 (Fig. 1 A and B, and SI Appendix Table S2). No significant differences were observed between leaves 1-2 and 3-4 (Fig. 1 B). We concluded that the increased resistance to *Pto* DC3000 was associated with vegetative phase change in Col-0.

**Figure 1.**
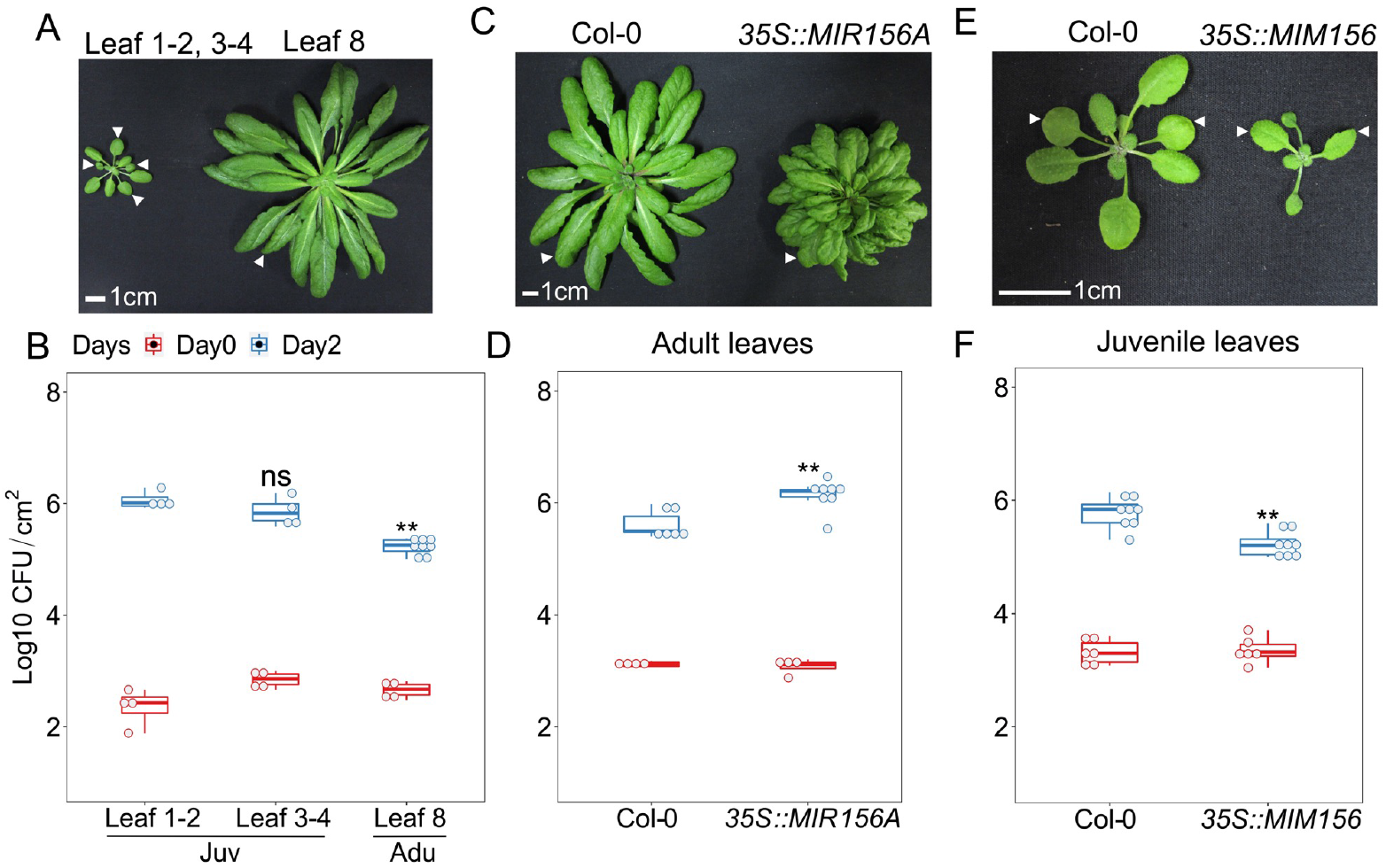
MiR156 suppressed age-related resistance to *Pto* DC3000 in *Arabidopsis*. **A**, developmental phenotype of a 4-week (left) and 7-week (right) Col-0 plant grown under short-day conditions. Arrows point to leaves 1,2,3,4 and leaf 8 in the left and right plants, respectively. **B**, *Pto* DC3000 bacterial growth in juvenile and adult leaves of Col-0. **C**, developmental phenotype of a 7-week Col-0 and *35S::MIR156A* plant. Arrows indicate an adult leaf of Col-0 or a leaf from a similar position in *35S::MIR156A*. **D**, bacterial growth in adult leaves of Col-0 and *35S::MIR156A*. **E**, developmental phenotype of a 4-week Col-0 and *35S::MIM156* plant. Arrows indicate leaves 1 and 2 on each plant. **F**, *Pto* DC3000 bacterial growth in juvenile Col-0 and *MIM156* leaves 1-2 on Day 0 and Day 2. Scale bar = 1 cm. Day 0, the day of *Pto* infection. Day 2, two days post-infection. CFU/cm^2^, bacterial colony forming unit per square centimeter of a leaf. Juv, juvenile leaves, Adu, adult leaves. The student t-test was used for statistical analysis. Each genotype was compared with Col-0, ns, not significant, *, p < 0.05, **, p < 0.01. The same annotation is used for bacterial growth dot-box plots shown in Figures 2 and 7. Repeats of bacterial growth are presented in SI Appendix, Table S2.

Since miR156 level in fully expanded leaves drops from the juvenile to adult transition (35), we hypothesized that a high level of miR156 suppresses immunity in juvenile stage. We compared bacterial growth in adult leaves (leaves ≥ 8) from Col-0 and leaves at the same position from transgenic plants overexpressing *MIR156A* under a constitutive 35S promoter from TMV (*35S::MIR156A*). Expressing *MIR156A* in adult leaves led to accelerated production of juvenile leaves, marking the prolonged juvenile phase as previously reported (20, 21) (Fig. 1 C). Interestingly, it also led to an increase in bacterial growth when compared with leaves at the same position in Col-0 (Fig. 1 D). Consistently, knocking down miR156 activity by a target mimicry, 35S::*MIM156* (36) (*MIM156*) displayed enhanced disease resistance (Fig. 1 E and F). These evidences suggests that high accumulation of miR156 suppresses resistance to *Pto* DC3000 in the juvenile phase.

### miR156-regulated SPL10 promotes ARR in adult phase

To test which miR156-targeted *SPL* contributes to ARR_VPC_, we first screened disease phenotype in the gain-of-function mutants among 9 available (out of 11) miR156-targeted SPLs (Fig. 2 A and SI Appendix, Table S2). We examined the disease phenotype in juvenile leaves expressing individual miR156-resistant *SPL* gene under its own native promoter (*rSPLs*) (24). Low levels of endogenous *SPL* transcripts in juvenile leaves provided a sensitized background to test the function of *rSPLs*. We found that leaves 1-2 from *rSPL2, 10, 9* and *13* showed increasing resistance compared to wild type, but *rSPL3, 4, 6, 11* and *15* did not change resistance to *Pto* DC3000 (Fig. 2 A and SI Appendix, Table S2 and Figure S2). Thus, a subset of miR156-regulated SPLs was sufficient to enhance immunity in juveniles.

**Figure 2.**
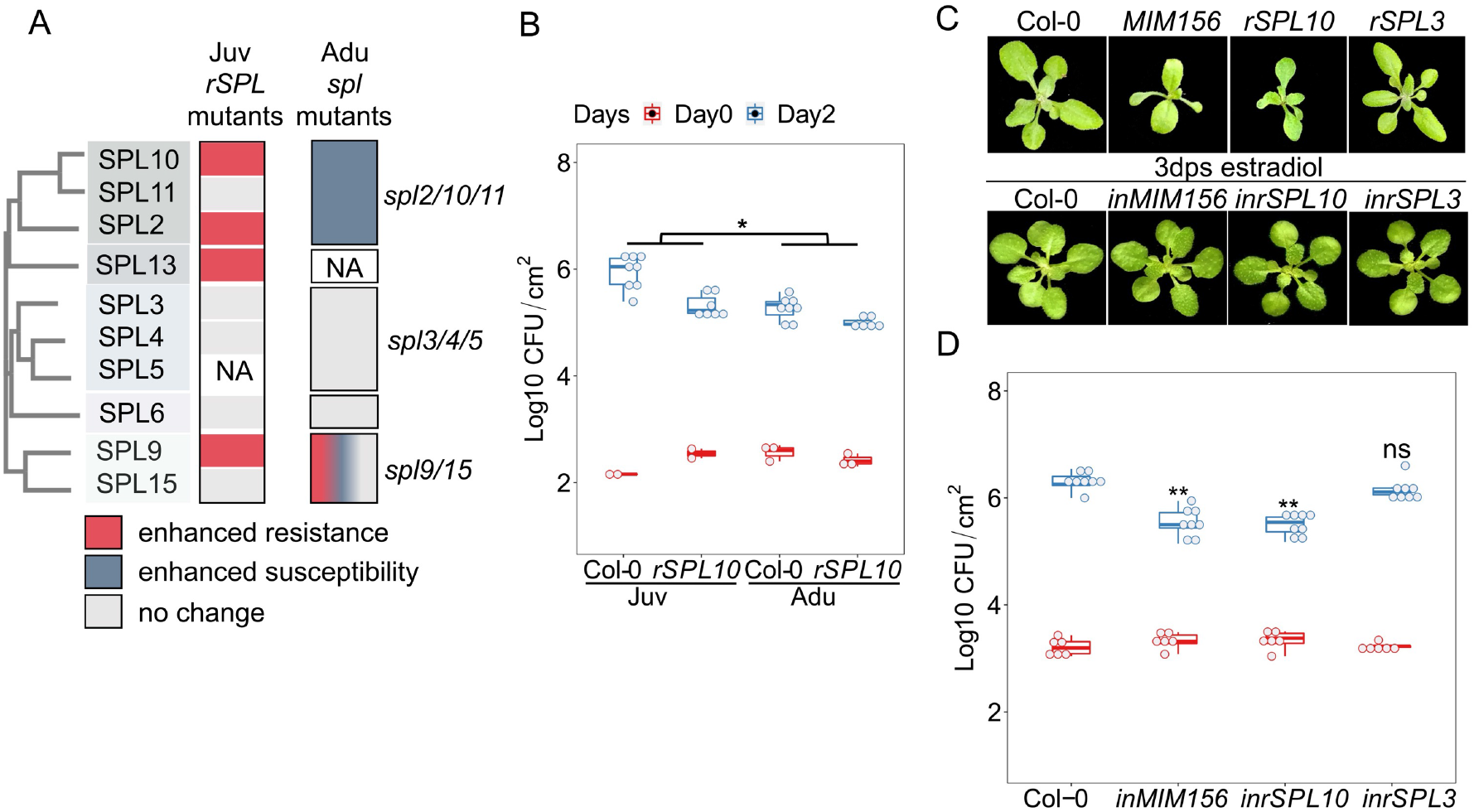
MiR156-targeted SPL10 promoted resistance to *Pto* DC3000. **A**, disease phenotype of *Pto* DC3000 in *SPL* gain and loss-of function mutants. NA, not available. **B**, bacterial growth in leaves 1-2 and leaves 11 from Col-0 and *rSPL10*. **C**, developmental phenotype of Col-0, and early phase change phenotypes of transgenic plants expressing stable *MIM156, rSPL10* and *rSPL3* on ½ MS plate (top panel); developmental phenotype of Col-0, *estradiol-inducible(in)MIM156, inrSPL10* and *inrSPL3* at 3 days after estradiol treatment (bottom panel). **D**, bacterial growth in leaves 1-2 from 4 -week-old Col-0, *inMIM156, inrSPL10* and *inrSPL3* on Day 0 and Day 3. Emmeans package in R was used for statistical analysis in 2 A and 2 B. The student t test was used for statistical analysis in 2 A and 2 C, and each genotype was compared with Col-0 wild type, ns, not significant, *, p < 0.05, **, p < 0.01. Repeats for the experiments here are shown in SI Appendix, Table S2.

To check the necessity of SPLs in adult conferred immunity, we tested disease phenotypes in adult leaves with loss of function *spl* combinatorial mutants based on amino acid sequence similarity (Fig. 2 A; SI Appendix, Table S2 and Figure S2). There are 5 phylogenic clades for miR156-targeted *SPL* genes, *SPL3/4/5, SPL6, SPL2/10/11, SPL9/15*, and *SPL13A/13B* (Fig. 2 A) (26, 37). In Col-0 background, *SPL10* and *SPL11* (78% amino acid identity) reside in a 1.6 kb tandem duplication (26). S*pl2/10/11* showed enhanced susceptibility to *Pto* DC3000 (Fig. 2 A). While the phenotypes of *spl2 and spl10* single mutants were wild type-like, *spl10/11* mildly reduced disease resistance (SI Appendix, Table S2), indicating that *SPL2, 10* and *11* redundantly contributed to immunity in adult leaves. Importantly, although *rSPL10* carried a higher resistance than wild type in juvenile leaves (Fig. 2 A and B), the difference between the two genotypes was much smaller in the adult stage (Fig. 2 B), supporting that SPL10 enhanced resistance is age dependent. *Spl3*/*4*/*5* triple or *spl6* single mutations did not alter resistance to *Pto* DC3000 (Fig. 2 A; SI Appendix, Table S2). The lack of disease phenotypes against *Pto* DC3000 in the gain- and loss-of-function mutants of *SPL6* is consistent with a previous report (38). *rSPL9* enhanced resistance, but the *spl9/15* double mutant displayed unstable phenotypes, which may be resulted from redundancy among other SPL members (Fig. 2 A and SI Appendix, Table S2). Altogether, the data confirmed the specialized function of SPLs in ARR_VPC_.

To further determine whether the disease phenotypes of *SPL10* mutants were pleiotropic effects caused by SPL10-mediated leaf morphogenesis, we assayed bacteria multiplication in transgenic plants expressing either estradiol inducible *rSPL3, rSPL10* or *MIM156* (Fig. 2 C). Transgenic plants growing on ½ MS medium supplemented with estradiol showed typically early phase change phenotypes, indicating that the transgenes were functional (24, 39). In line with the data above (Fig. 2A), applying estradiol 12 hrs before inoculation suppressed bacterial multiplication in juvenile leaves of inducible *rSPL10* and inducible *MIM156* lines, but not for that from inducible *rSPL3* plants (Fig. 2 D). Notably, the leaf morphology was comparable between wild type and estradiol induced plants (Fig. 2 C), indicating that miR156-SPL10 controlled disease resistance and leaf morphology can be decoupled.

### The basal expression of defense genes is high in the adult stage

To explore the transcriptional signature of ARR_VPC_, we performed RNA sequencing to compare juvenile (leaves 1-2) and adult (leaves ≥ 8) transcriptomes at 3 hours after mock treatment or *Pto* DC3000 infection (Fig. 3 A). An early time point was chosen because change of chromatin accessibility could be detected at 3 hrs post *Pto* infection (40). Under mock treatment, we identified 2002 and 2320 genes that were respectively up- or down-regulated in adult leaves compared with juvenile leaves, hereafter Adu_no infection (nof)_ (LFC = 0.58, padj < 0.05) (Fig. 3 B, and SI Appendix, Table S3 and Figure S3 C and S3 D). As expected, *SPL3/4/5* were up-regulated in adult samples (SI Appendix, Table S3). In addition, the Gene Ontology (GO) enrichment analysis revealed that vegetative phase change related GOs, such as adaxial/abaxial axis specification (41) were enriched in the Adu_nof_ DEGs (Fig. 3 D and SI Appendix, Table S6.1), confirming that our juvenile and adult samples represented two distinct developmental phases (SI Appendix, Fig S3 A-B and SI Appendix, Table S3). Intriguingly, we observed that immunity related GOs were also enriched in the up-regulated Adu_nof_ DEGs, including defense response to bacterium/fungus and response to SA (Fig. 3 D). Here, jasmonic acid (JA) signaling-mediated defense was downregulated, coinciding with the antagonistic interaction between JA and SA in plant immunity (42). The enrichment of pro-defense genes in up-regulated Adu_nof_ indicated that adult plants had primed defense signaling.

**Figure 3.**
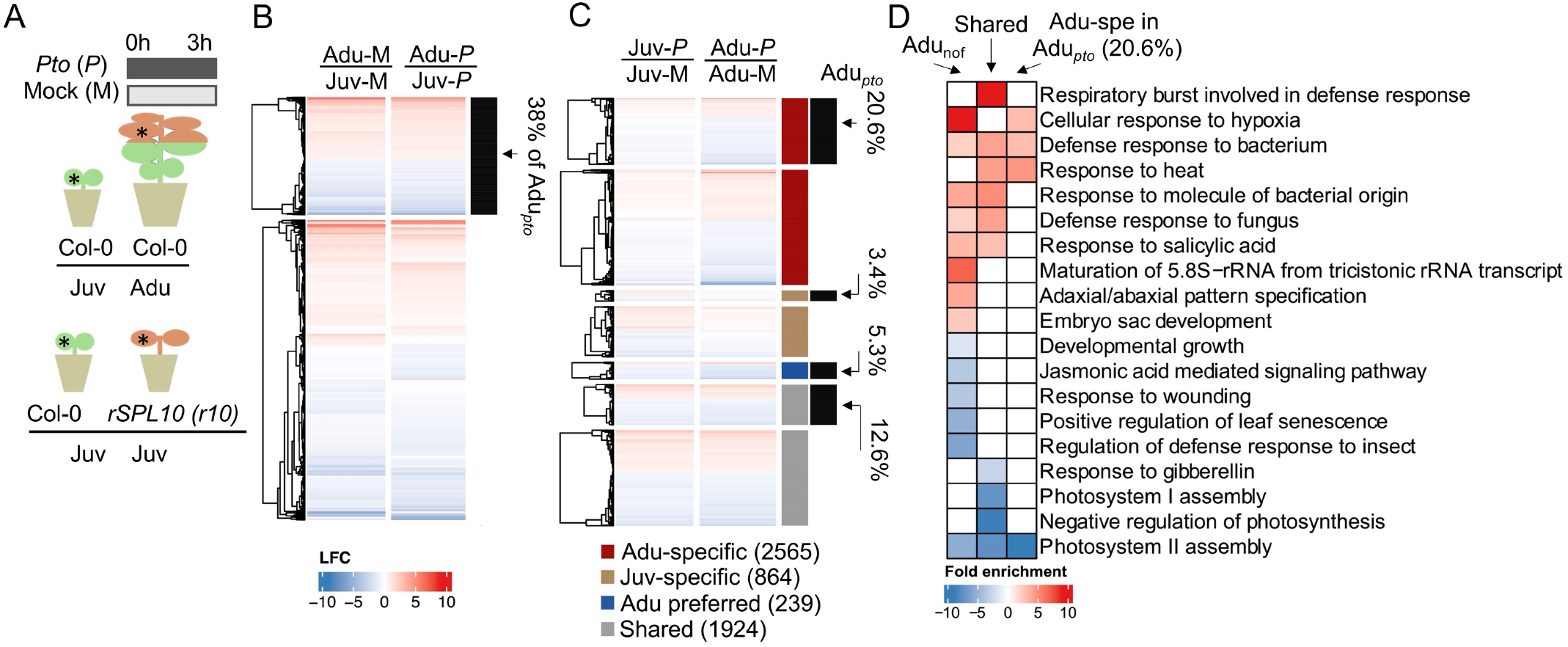
Basal transcript level of defense genes were heightened over vegetative phase change. **A**, experimental settings of RNA-seq. Non-infected state and challenged (3 hours after *Pto* inoculation) transcriptomes from leaf 1-2 (Col-0 and *rSPL10*) and leaf 11 (Col-0) were compared. * indicates examples of leaf samples that were collected for RNA-seq. Green: juvenile leaves; brown: adult leaves. **B**, an expression profiling of DEGs identified in mock-treated adults against mock-treated juvenile leaves. DEGs, differently expressed genes with LFC ≥ ±0.58 and padj ≤ 0.05. LFC, log2 fold change of gene expression. Padj, adjusted p value. Color-codes in the heatmap, blue is for down-regulated DEGs and red is for up-regulated DEGs. Euclidean distance was used for calculating distance. Complete linkage was applied to structure the hierarchical clustering. Expressing profiles of DEGs derived from Adu-M/Juv-M are mapped in the first column of the map. The expressing profiles of those DEGs in Adu-*P*/Juv-*P* are shown in the second column of the map. Genes that were DEGs in both Adu-M/Juv-M and Adu-*P*/Juv-*P* are marked by the black bar on the right. 38% indicates the percentage of those overlapping DEGs in Adu_*pto*_. **C**, a profile of *Pto*-triggered DEGs came from Juv-*P*/Juv-M and Adu-*P*/Adu-M. Adult-specifically *Pto*-triggered DEGs (dark red), Juvenile-specifically triggered (light brown), adult preferentially triggered (deep blue), and commonly triggered in both adults and juveniles, i.e., shared (light grey) are marked by the first column of bars on the right. The percentages (20.6%, 3.4%, 5.3%, 12.6%) and corresponding black bars indicate the proportion of each DEG category that overlaps with Adu_*pto*_. **D**, representative GO terms enriched in DEGs of Adu_nof_, DEGs triggered by *Pto* DC3000 in both adult and juvenile leaves (the Shared) and DEGs specifically triggered by *Pto* in adult phase within the total Adu_*pto*_ (20.6%). Red and blue color blocks refer to GOs enriched in up- and down-regulated DEGs, respectively. Only GO terms with FDR < 0.05 were deemed as enriched here. Fold enrichment was based on hypergeometric tests within the range of the DEG set used for each GO analysis relative to *Arabidopsis* genome. The analysis was done using the TAIR Gene ontology website (geneontology.org). The same GO analysis and color-coding are used for Figure 4 C.

For defense induction state, we investigated genes that were differentially induced/suppressed by infection in juvenile and adult stages. *Pto* treatment triggered 3027 and 4728 DEGs in juvenile and adult leaves, respectively (SI Appendix, Table S3-S4 and Fig S3 C and S3 D). Among the 2163 *Pto-*triggered DEGs shared between juvenile and adult leaves (2163 = 239 + 1924 in Fig. 3 C and SI Appendix, Table S3-S4 and Fig S3 C and S3 D), the shared up-regulated DEGs were enriched with well-characterized defense responses such as respiratory burst, defense response to bacterium and response to salicylic acid (Fig. 3 D). Meanwhile, photosynthesis and light harvesting signaling pathways were enriched in down-regulated DEGs, indicating that a pathogen-induced transcriptomic reprogramming switched from development to defense regardless of plant age (Fig. 3 D, SI Appendix, Table S6.1). A total of 864 DEGs were only induced/repressed by *Pto* in the juvenile stage, and there are 2565 DEGs were adult-specific (Fig. 3 C). Cutin biosynthetic and wax biosynthetic processes were enriched in adult-specific *Pto*-induced DEGs, consistent with a previously suggested consolidation of constitutive defense in ARR (43) (SI Appendix, Table S6.2).

We noticed that a portion of genes specifically induced in the adult stage did not eventually have higher transcription level than those in juvenile stage. It is arguable whether inducibility of a gene alone contributes to ARR. So, we further searched for genes whose absolute expression levels were different between juvenile and adult leaves after *Pto* DC3000 treatment (hereafter Adu_*pto*_). Genes that were specifically induced/repressed by infection in the adult stage count for 20.6% (934 out of 4528, SI Appendix, Table S3) of the Adu_*pto*_ (Fig. 3 C). There, cellular response to hypoxia, defense response to bacterium and response to heat were over-represented, indicating an age-dependent *Pto* response that may cope with abiotic stresses (Fig. 3 D). A 5.3% (239 out of 4528, SI Appendix, Table S3) of Adu_*pto*_ were also triggered by *Pto* in the juvenile stage, but the amplitude of change was preferentially higher in the adult stage (Fig. 3 C). Thus, ARR_VPC_ strengthened a sector of juvenile-defense regulon as well as activated adult-specific defense genes. In addition, 38% (1722 out of 4528) of Adu_*pto*_ were not *Pto*-triggered but already had differential expression between juvenile and adult leaves before infection (Fig. 3 B, SI Appendix, Table S3). Of the 38%, GOs pertaining to defense, such as SA signaling, together with adult-related developmental pathways were enriched (Fig. 3 D, SI Appendix, Table S6.1). Finally, 20.1% Adu_*Pto*_ (910 out of 4528) was not induced either by infection or age alone. These DEGs could be resulted from a synergistic interaction between age and infection (Fig. S3 C and SI Appendix, Table S3). Taken together, we discovered that ARR transcriptome changes could contribute to the elevation of defense signaling at non-infected state, the adult-specific inducible defense as well as the strengthened juvenile defense.

### Overexpressing SPL10 recapitulates the ARR transcriptomic landscape in juvenile leaves

To delineate the contribution of SPL10 to ARR at the transcriptomic level, we investigated the *rSPL10 (r10)* induced DEGs at non-infected and *Pto*-infected states (Fig. 3 A). A total of 3211 genes (1859 up-regulated and 1352 down-regulated) were differentially expressed in leaves 1-2 between Col-0 and *r10* at non-infected state (*r10*_nof_). Among them, 936 out of 3211 *r10*_nof_ DEGs were co-regulated in adult leaves, attributing to 21.7% (936/4322) of the Adu_nof_, being consistent with the function of SPL10 in specifying adult fate (Fig. 4 A, SI Appendix, Table S5). GO terms associated with immune signaling including systemic acquired resistance were enriched in the 936 co-upregulated DEGs between *r10*_nof_ and Adu_nof_ (Adu/*r10*_nof_) (Fig. 4 C, SI Appendix, Table S6.3). After bacterial infection, we identified 2621 DEGs between leaves 1&2 from *r10* and Col-0. 604 of those DEGs were also Adu_*pto*_, occupying 13.3% (604/4528) of total Adu_*pto*_ (Adu/*r10*_*Pto*_) (Fig. 4 B, SI Appendix, Table S5). Defense-related GOs were enriched in those co-regulated DEGs (Fig. 4 C, SI Appendix, Table S6.3). The observations support that SPL10 activated a sector of adult immune response to *Pto* DC3000.

**Figure 4.**
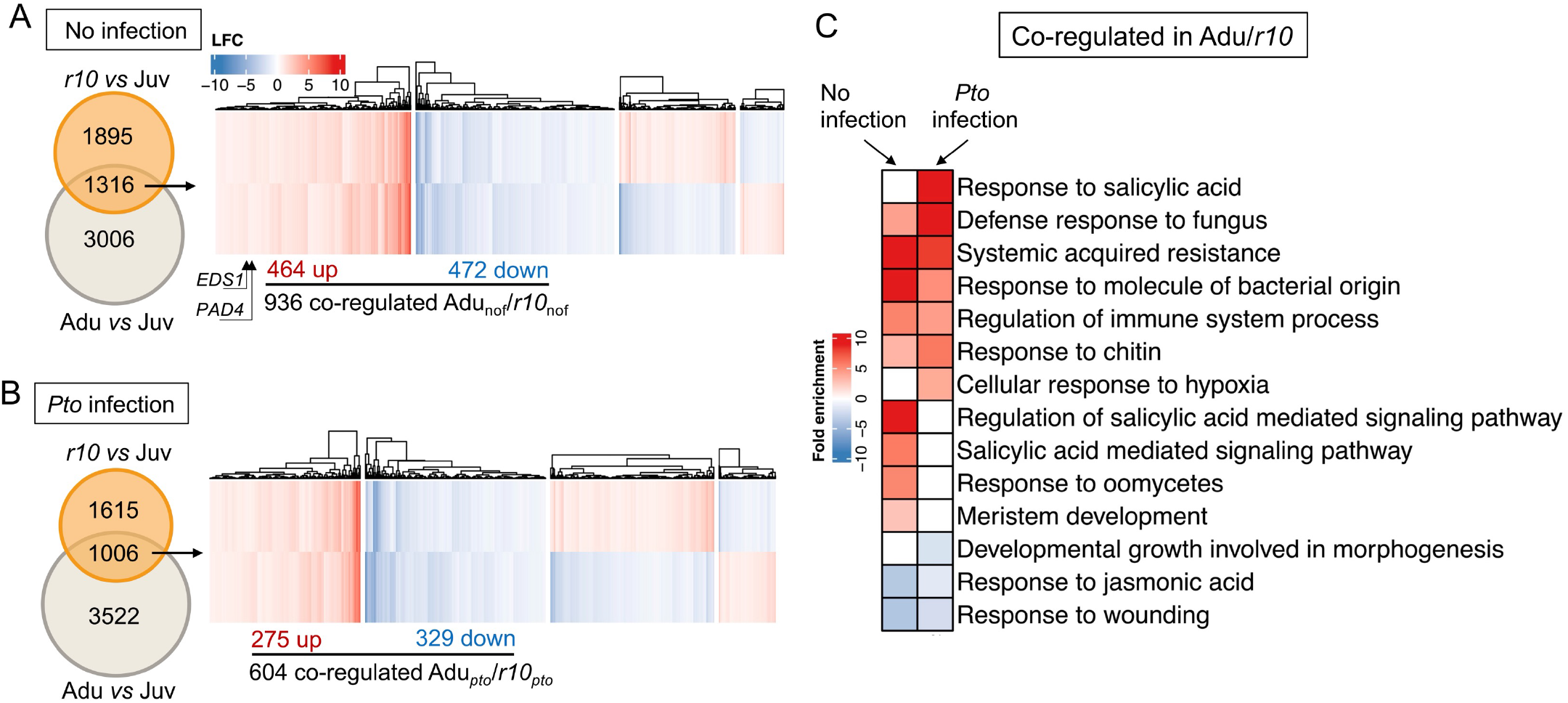
*rSPL10* transcriptomes resembled that of adult leaves. **A**, an expression profiling of co-regulated DEGs by adult and *rSPL10* leaves at non-infected state compared with juvenile leaves. *EDS1* and *PAD4* were identified as Adu/*r10* co-upregulated DEGs under non-infected state, which are indicated in the heatmap with arrows. **B**, expressing profiles of co-regulated DEGs by adult and *rSPL10* from Adu-*P* or *r10*-*P* against Juv-*P*. For the clustered heatmaps in 4 A and B, blue represents down-regulated DEGs and red is for up-regulated DEGs. Euclidean distance was used for calculating distance in the partition around medoids (PAM) clustering (k=4). **C**, GO enrichment of co-regulated adult and *r10 vs* juvenile DEGs in non-infected and *Pto*-infection states.

### Disrupting SA signaling compromised the SPL10-mediated ARR

Since SA-related GO terms were enriched in the co-upregulated Adu/*r10*_nof_ (Fig. 4 C), we first measured the age- and SPL10-effects on SA signaling. We assembled a core SA regulon by overlapping DEGs induced by SA and its synthetic inducer benzothiadiazole (BTH) (44, 45) (Fig. 5 A). 81.3% of the core SA regulon (527 activated and 239 repressed genes) were detected in our sequencing data. SA-activated (199/527) and -repressed (50/239) markers were enriched in Adu_nof_ (Fig. 5 B and SI Appendix, Information S6). SA-activated genes were also enriched in *r10*_nof_ (Fig 5 B and SI Appendix, Information S6), denoting that SPL10 contributed to enhancing SA signaling in the adult phase. Consistently, accumulation of Salicylic acid beta-glucoside (SAG) (an inactive SA form, stored in vacuole) was higher in adult than that in juvenile leaves (Fig. 5 C and SI Appendix, Table S2). The accumulation of free SA showed a similar trend (Fig. 5 C and SI Appendix, Table S2). Next, we sought to validate whether elevated SA response in adult phase depended on the temporal expression of *SPL10*. We selected four age-dependent SA-activated genes, *AT3G60470, BG3, ATLTP4*.*4* and *PR5*. Their expressions were compromised in the adult leaves of *spl2/10/11* (Fig. 5 D). Noticeably, the upregulation of the four SA responsive genes were also observed in adult or *r10* leaves when plants were grown on sterile plates (Fig. 5 E). In conclusion, the age-related increase of SA response in leaves required *SPL10*, and was not primed by pre-exposure to microbes. TheEDS1-PAD4 protein complex is essential for SA-mediated defense signaling (46). We found that both genes were upregulated in adults and *r10* (Fig. 4 A). Furthermore, a significant overlap was found between the EDS1-PAD4 core regulon (EP-core, (46) and genes co-upregulated by adults and *r10* (Fig. 5 F), implying that the EDS1-PAD4 mediated SA signaling pathway could be differentially activated in juvenile and adult stage due to the temporal expression of SPL10.

**Figure 5.**
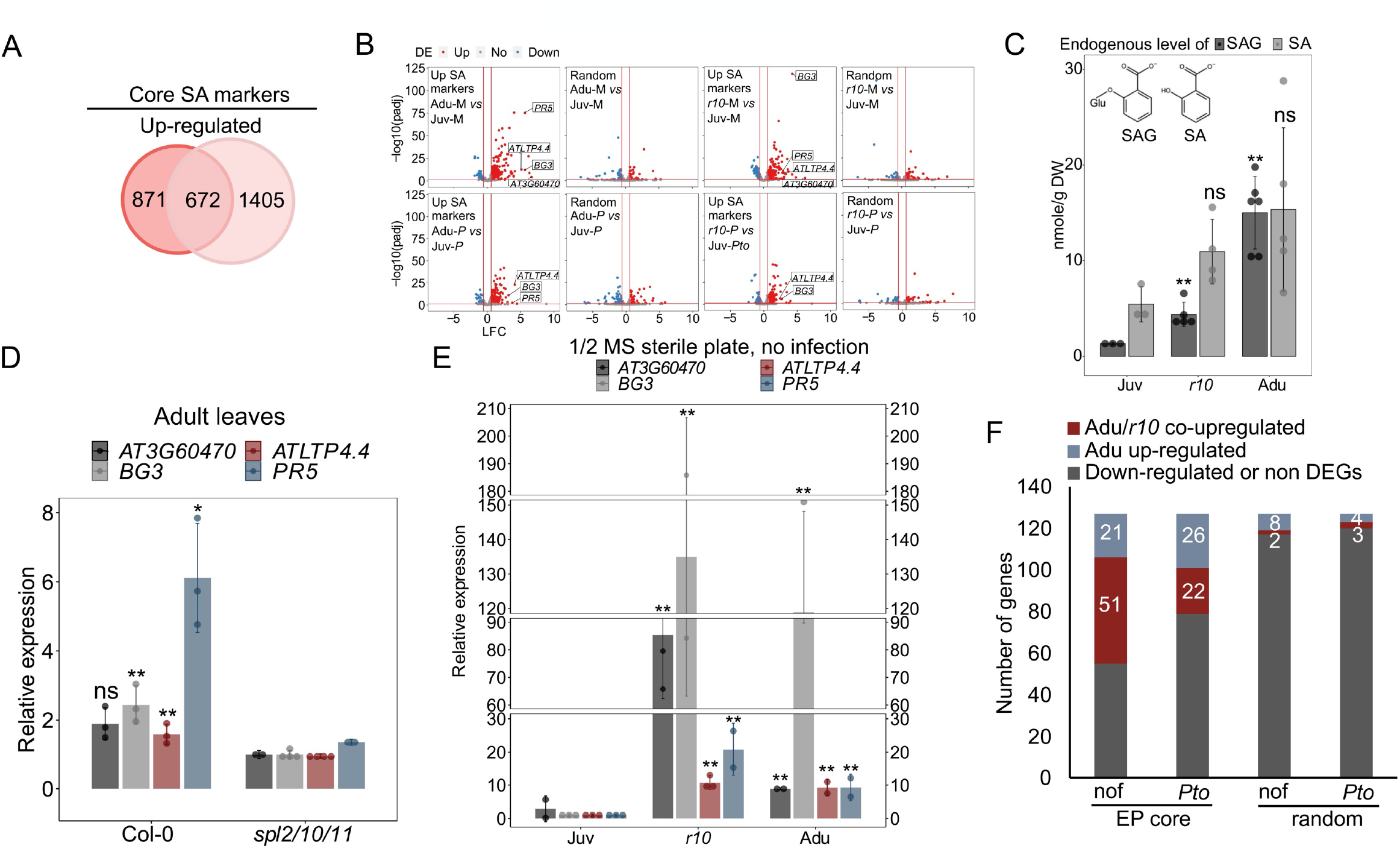
Salicylic acid (SA) biosynthesis and signaling were enhanced by SPL10 in adult phase. **A**, 672 up-regulated core SA markers were defined via overlapping DEGs pools from Ding Y *et al*., 2018 (dark circle) and Yang L *et al*., 2017 (light circle). **B**, expression patterns of 527 detected (out of 672) up-regulated core SA markers in adult and *r10* leaves, under non-infected and *Pto* infection states. Four random pools of detected genes (450 genes per set) derived from each pair-wise comparison exhibits here were chosen as negative controls. Up, upregulated. No, no change. Down, downregulated. Y axis shows the negative logarithm transformed adjusted p value, -log10(padj). X axis displays the value of Log2 fold change (LFC) of gene expressions. M, mock. *P, Pto* DC3000. P < 0.0001 (Hypergeometric test, done by GeneOverlap R package) was reproducibly output from each overlap between core upregulated SA markers and Adu_nof_, Adu_*pto*_, *r10*_nof_ and *r10*_*pto*_ (SI Appendix, information S6). The P values for 3 out of 4 random controls were not significant (SI Appendix, information S6). **C**, endogenous SAG and free SA accumulation in juvenile, *r10* and adult leaves at non-infected state. DW, dry weight. Repeats for the experiment are shown in SI Appendix, Table S2. **D**, age-associated expression of four SA markers in adult leaves from Col-0 and in comparable leaves of *spl2/10/11*. Similar results were seen two times. **E**, the qPCR of the four SA marker genes in juvenile, *r10* and adult leaves on sterile 1/2 MS plates. Similar results were seen three times. **F**, overrepresentation of EDS1-PAD4 core regulon (EP core) in Adu/*r10*_nof_ and Adu/*r10*_*pto*_. The EP core markers were defined in Cui *et al*., 2017 (46). 127/155 of the EP core markers were detected in this work. Randomly selected 127 genes from our RNA-seq dataset were used as controls.

Out of the 936 DEGs co-regulated by adult stage and *rSPL10* (Fig. 4 A), 400 of them were associated with SPL10-binding sites identified in a ChIP-seq experiment of SPL10 by Ye *et al* (Fig. 6 A) (25), indicating that these genes are likely direct targets of SPL10. A de novo motif discovery algorism identified potential SPL-binding sites in 203 of the co-up-regulated DEGs (Fig. 6 A), but not in the 197 down-regulated genes. In addition, experimentally validated SPL TF binding sites (47) were enriched in the promoter of the 203 genes (SI Appendix, Fig S5 A). Interestingly, the promoter region and gene body of *PAD4* contains two potential SPL10-binding GTAC-containing motifs (Fig. 6 B) (25). We generated a genomic reporter line of SPL10 (*proSPL10::rSPL10-YFP*). The transgenic line showed similar early phase change phenotypes observed in the *proSPL10::rSPL10-GUS* line, indicating that the fusion had a normal SPL10 function (SI Appendix, Fig S5 B). Using ChIP-qPCR, we validated that the motif 1 (M1, 186-182 bp away from the TSS), but not motif 2 was associated with SPL10 at uninfected state (Fig. 6 B). Furthermore, the transcript level of *PAD4* was reduced only in the *spl2/10/11* adult leaves but not in the juvenile leaves (Fig. 6 C), indicating that the temporal expression of *PAD4* depends on SPL10. Taken together, these observations suggest that SPL10 directly promotes *PAD4* expression in the adult stage.

**Figure 6.**
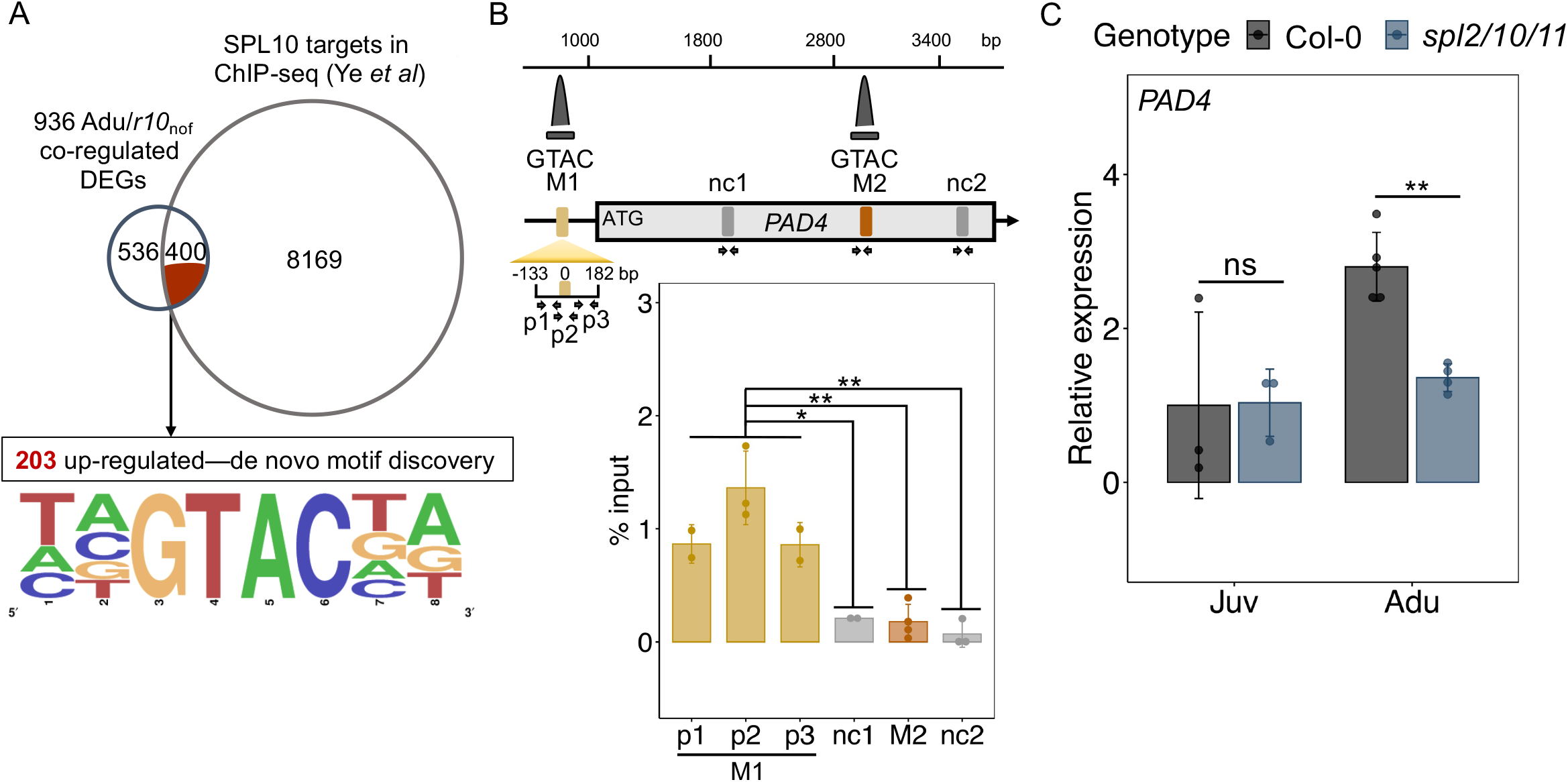
*PAD4* was a direct target of SPL10. **A**, Overlap between Adu/*r10*_*nof*_co-regulated genes and potential SPL10 targets defined according to ChIP-seq data from *Ye et al*. (upper panel). A motif discovery and enrichment analysis of the 203 Adu/*r10*_nof_ co-upregulated DEGs (lower panel). **B**, SPL10 bound to a GTAC-containing motif upstream of *PAD4*. qPCR following chromatin immunoprecipitation of *rSPL10*-YFP for motifs (M1 and M2) and negative control sites (nc1 and nc2) are at least 600 bp away from M1 and M2 at the *PAD4* genomic region. Three primer sets (p1-p3) were used to amplify the M1 site. Relative locations of ChIP peaks (dark grey) derived from *Ye et al*, primers (arrow pairs) and the tested sites (color blocks) were indicated in the schematic diagram. The student t test was performed to compare and indicate the significance of difference between the sites. **C**, qPCR of *PAD4* transcripts level in juvenile (Juv) and adult (Adu) leaves of Col-0 and *spl2/10/11*. The student t test, ns, not significant, *, p < 0.05, **, p < 0.01.

To probe the genetic interactions of SA signaling and the miR156-SPL10 mediated ARR, we first tested the ARR_VPC_ phenotype in the mutant of *SALICYLIC ACID INDUCTION DEFICIENT 2* (*sid2-1*, defective in pathogen-induced SA biosynthesis) and *NONEXPRESSER OF PR GENES 1* (*npr1-1*, defective in SA perception) mutants. Neither of those mutants altered the timing of vegetative phase change. As expected, both mutants in adult stage were more susceptible than Col-0 (Fig. 7 A). The difference of bacterial growth between wild type and the mutants in juvenile stage was less pronounced (Fig. 7 A), in agreement with the age-dependency of SA-mediated disease resistance. We then introduced *sid2-1* mutation into *MIM156* background (SI Appendix, Fig. S5 A). Although precocious morphological traits were shared between *MIM156* and *MIM156/sid2-1*, the bacteria level in *MIM156/sid2-1* phenocopied that of *sid2-1* (Fig. 7 B), suggesting that *SID2* acts downstream of miR156. Similarly, loss of function mutation of *EDS1* reversed the enhanced disease resistance phenotype in *r10* plants (Fig. 7 C, SI Appendix, Fig. S5 B). The leaf morphology phenotype of *r10* plants was not changed by *eds1*.*2*, which further confirmed that age-associated leaf morphology and disease resistance can be decoupled (Fig. 7 C). Consistent with the molecular evidence that SPL10 binds to the promoter of *PAD4*, the ARR_VPC_ phenotype was compromised in *pad4-1* and *eds1*.*2* mutant (Fig. 7 D). In essence, miR156-SPL10 promoted resistance through *SID2* and *EDS1-PAD4*-dependent SA signaling.

**Figure 7.**
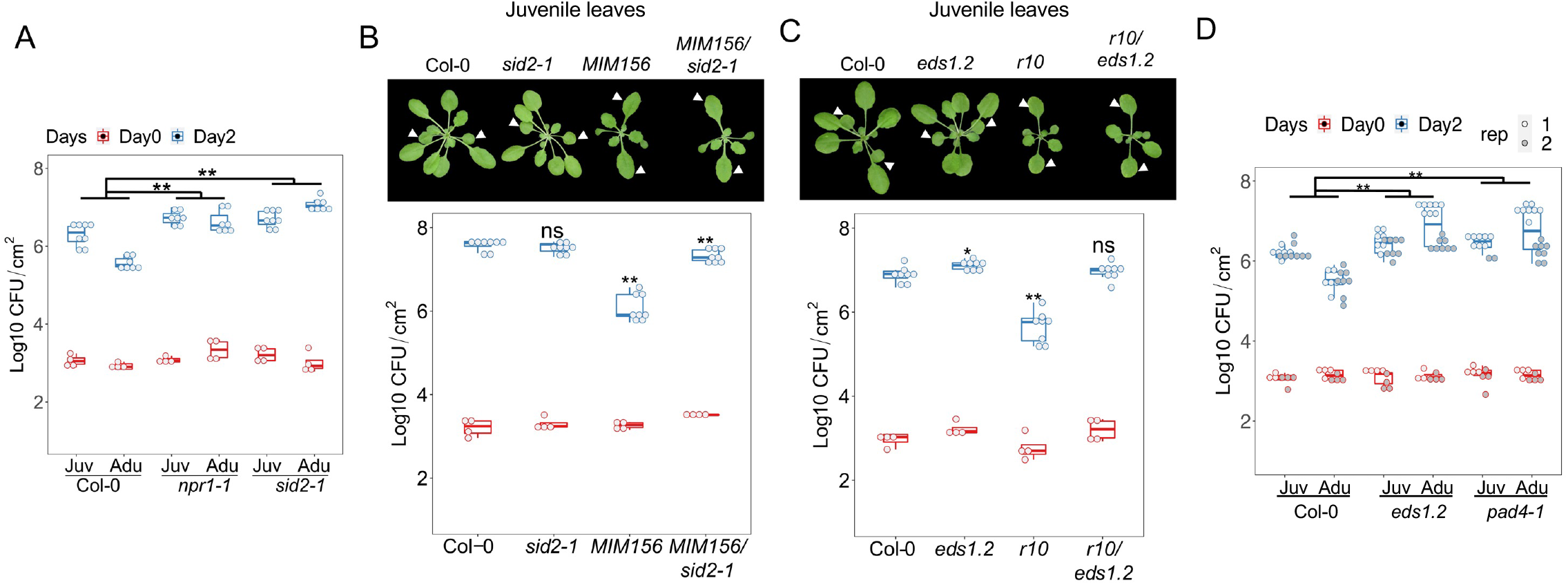
SA biosynthesis and signaling components were required for the SPL10 regulated ARR_VPC_. **A**, a comparison of *Pto* DC3000 growth between the juvenile and adult leaves (ARR phenotype) of Col-0, *sid2-1* and *npr1-1*. **B**, developmental phenotype (top) and the bacterial growth (bottom) in 4-week-old leaves 1 and 2 from Col-0, *sid2-1, MIM156* and *MIM156/sid2-1*. **C**, developmental phenotype of 4-week-old Col-0, *eds1*.*2, r10* and *r10 /eds1*.*2*. Note the similar leaf shape between *r10* and *r10*/*eds1*.*2;* (bottom panel) bacterial multiplication in leaves 1 and 2 derived from Col-0, *eds1*.*2, r10* and *rSPL10/eds1*.*2*. Arrows indicate leaves 1-2. **D**, the ARR phenotyping of *eds1*.*2* and *pad4-1* infected with *Pto* DC3000. Emmeans package in R was used for statistical analysis in 7 A and D. The student t test was used for statistical analysis in 7 B-C, and each genotype was compared with Col-0 wild type, ns, not significant, *, p < 0.05, **, p < 0.01.

## Discussion

In this research, we uncovered an ARR mechanism regulated by the miR156-SPL pathway, linking immune maturation to an intrinsic aging mechanism (SI Appendix, Fig. S6). In plants, ARR occurs during predicable developmental transitions that are often accompanied by morphological changes. Our research provides a potential mechanism for the temporally coordinated change of immunity and morphogenesis, where the same molecular clock, miR156, controls different *SPL* genes to specify immune and developmental traits. For instance, SPL2/10/11, but not SPL3/4/5/6, are required for resistance against *Pto* DC3000 (Fig. 2 A). Interestingly, NbSPL6 is required in the N-mediated TMV resistance in tobacco (38). The homologue of *NbSPL6* in *Arabidopsis, AtSPL6*, is necessary to the full ETI response triggered by *Pseudomonas syringae* effector AvrRps4, but not basal resistance to *Pto* DC3000 (38) (Fig. 2 A). It is possible that the broad spectrum of resistance associated with ARR requires multiple SPL family members to activate different defense pathways. As rSPL10 only influenced 21.7% of Adu_nof_ and 13.3% of Adu_*pto*_, other SPLs may contribute to the rest of adult defense to *Pto* and/or to other pathogens (Fig. 4 A and 4 B). Alternatively, the coordinated maturation of development and immune system may be achieved by functional switch of the same SPL protein. In rice plants, *Ideal plant architecture 1* (*IPA1*)/*OsSPL14* and *OsSPL7* promote growth in a non-infection status. It binds to an alternative cis-regulatory element to activate defense when challenged by the bacterial pathogen *Xanthomonas oryzae pv. Oryzae* (*Xoo*) (48, 49). In *Nicotiana benthamiana* and tomato plants, temporal reduction of miR6019/6020 allows its target, *N* gene, to mediate age-dependent resistance to tobacco mosaic virus (TMV) (50). The temporal expression pattern of miR6019/6020 mimics that of miR156 in tobacco. It would be interesting to explore the genetic relationship between those two miRNAs in regulating ARR.

We showed that defense GO terms were enriched in up-regulated *rSPL10*_nof_ and Adu_nof_. Among those, components of SA pathway, such as *SID2* and *NPR1* were required for ARR_VPC_ (Fig. 6). We observed the up-regulation of SA response accompanied by a reduction of JA response in both Col-0 adults and the *rSPL10* line (Fig. 3 and 4). JA and SA often act antagonistically to fine-tune immune response to multiple pathogens. Our ARR transcriptome results showed that the antagonism also occurred in an age-dependent manner (Fig 3 C). Upregulation of SA production and response have also been observed during vegetative-floral transition in both tomato and tobacco plants (51). In *Arabidopsis, SVP* and *SOC1* genes regulating floral induction also transcriptionally promote age-dependent increase of SA, independent from flowering traits (52, 53). We did not observe differential expression of *SVP* or *SOC1* in juvenile *vs* adult transcriptome, suggesting the *SVP-SOC1* module may not act in the ARR_VPC_ of *Arabidopsis*. On the other hand, we found that ARR_VPC_ was *NPR1*-dependent (Fig. 6 A), which is different from an age- and SA-associated but *NPR1*-independent gain of resistance observed before (54). It is likely that there are parallel aging pathways strengthen SA signaling during plant maturation.

In co-upregulated Adu/*r10*_*pto*_ DEGs, we noticed that cellular response to hypoxia was enriched (Fig. 4 D). Response to hypoxia of plants has been reported for counteracting submergence and waterlogging stress (55). Respiratory burst, as part of immune responses, is oxygen dependent. At the site of *Botrytis cinerea* infection, hypoxic response was induced, leading to the stabilization of subgroup VII of ETHYLENE RESPONSE FACTOR (ERF-VII) (46). Members in the ERF-VII can activate defense gene as well as hypoxic response (46). SPL10 may upregulate genes that react to low-oxygen environment when reactive oxygen species accumulates or respiration increases. How hypoxia response contributes to ARR against biotrophs and necrotrophs remains to be seen.

Plant innate immunity shares similarities to human innate immunity, ranging from structures and functionality of innate immune receptors and their downstream signaling cascades (56, 57). Our discovery of miR156-SPL signaling in ARR_VPC_ provides further evidence that heterochronic miRNAs coordinate aging and immunity *in planta*. In *Caenorhabditis elegans, let-7* family microRNAs are well-characterized regulators of developmental timing by specifying stage-specific cell fates in the hypodermal seam cell lineages (58, 59). Interestingly, let*-7-fam* miRNAs also repress the worm’s resistance to *Pseudomonas aeruginosa*, an opportunistic human pathogen (60). The dual function of *let-7-fam* in developmental timing and immunity is fulfilled through integrating downstream heterochronic genes and the p38 MAPK pathway (60). Thus, deploying heterochronic microRNAs pathway can be a cross-kingdom strategy to integrate immunity and developmental timing. It will be exciting to further dissect the genetic components and regulatory architecture of coordinated maturation of immunity and development.

## Material and methods

### Plant material and growth conditions

*Arabidopsis* wild type, transgenic lines and mutants used in this study were in a Columbia-0 (Col-0) genetic background unless mentioned otherwise. The genetic cross of *MIM156/sid2-1* and *r10/eds1*.*2* were generated from this research and progenies from F3 or F4 generation were used for phenotypic test. Information for mutants and transgenic lines can be found in SI Appendix, Table S1. Juvenile leaves were fully expanded leaves 1-2, or 3-4 from soil-grown 5-week-old plants. Adult leaves were fully expanded leaves that derived from 7-week-old plants. The adult phase of a leaf was confirmed by appearance of abaxial trichomes (41). Plants were sown on Fafard #3 Mix propagation soil. The planted seeds were then placed under 4 °C for 2 days and transferred to a growth room under 23°C/19°C day/night and with 45% humidity. Nine hours light and 15 hours dark photoperiod with 180 μmol m^-2^s^-1^ was used as short-day condition. Lighting was made through a 5:3 combination of white (USHIO F32T8/741) and red-enriched (interlectric F32/T8/WS Gro-Lite) fluorescent lights. Plant age was counted from the first day when seeds transferred to the growth room. Only plants used for Fig 5 E were grown on ½ MS plates in 24h with continuous light.

### Sampling strategy for studying ARR_VPC_

In our short-day growth condition, plants produced 50-60 leaves before bolting. The ontogenic age of a leaf influenced defense gene expression (SI Appendix, Fig. S1 A). To minimize the influence of ontogenic age of individual leaves (SI Appendix, Fig. S1 B), we measured the expansion rate of juvenile and adult leaves and harvested fully expanded juvenile and adult leaves from plants of different ages (SI Appendix, Fig. S1 C). Fully expanded leaves 1-4 derived from a 4- or 5-weeks old plants were sampled as juvenile leaves; fully expanded leaves (range from 8-13 depending on variations in plants) from 7-weeks old plants were sampled as adult leaves. Adult leaves showed characteristic abaxial trichome(s), blade serration and elongated petiole (21). Plants for adult samples were planted 2-3 weeks earlier than those for juvenile leaves. Juvenile and adult samples were collected at the same time for disease assay and transcriptome analysis.

### Bacterial growth assay

*Pto* DC3000 strain was grown under 28°C on King’s B solid medium (40 g/L proteose, 20 g/L glycerol and 15 g/L agar). The medium contained rifamycin for selection and cycloheximide to inhibit fungi growth. Glycerol stock of the bacterial strains stored under -80°C. Bacterial stock was streaked on plate for a 2-day growth and was re-streaked one day before inoculation. For infiltration, bacteria were collected from the plate and suspended in 10 mM MgCl_2_ solution. Bacterial suspension with concentration of 1 × 10^5^ CFU/mL was infiltrated in *Arabidopsis* leaves with a needless syringe. After inoculation, plants were covered by transparent lids for one hour. Day 0 samples were collected immediately after removing lids. Each sample contained four leaf discs that were derived from four individual leaves. Leaf samples were collected using the corer (the same size for all plants) and ground with homogenizer (OMNI International) and diluted serially. KB plates with 10 µL of bacteria suspension per sample were placed under room temperature for 2 days. Colony forming units were counted manually and normalized according to inside area of the corer. Day 2 samples were collected as described above two days post-infiltration (dpi).

### RNA sequencing and analysis

*Pto* DC3000 (1 × 10^8^ CFU/mL suspended in 10mM MgCl_2_) was infiltrated into juvenile and adult wild type (Col-0) leaves and leaves 1-2 from *rSPL10* plants. 10 mM MgCl_2_ was used as the mock treatment. Three hours after inoculation, 20 leaf discs with comparable size from 5-10 individual plants were cored and collected as one biological repeat per genotype per treatment. Three biological repeats were prepared for each genotype/treatment. For RNA isolation, plant tissues were flash-frozen in liquid nitrogen and then ground to fine powders using homogenizer (OMNI International). Total RNAs were extracted using E.Z.N.A. Total RNA kit (Omega BIO-TEK). RNA quality was assessed with a 2100 Bioanalyzer instrument (Agilent, RIN score ≥ 7, 28S/18S ≥ 1). RNA concentration was measured using a Nanodrop spectrophotometer (Thermo Scientific, RNA concentration ≥ 50 ng/μL, 260/280 ∼2.0). RNA samples were sequenced at BGI San Jose lab. Oligo dT based mRNA enrichment was followed by random N6 primer based reverse transcription. The synthesized DNA nanoballs were then sent for strand-specific mRNA sequencing (PE100) on a DNA Nanoball Sequencing (DNBseq) platform. The data was filtered using SOAPnuke software in BGI. ∼48 M clean reads with average Q30 ≥ 88.81% per sample were obtained.

Files of RNA-seq raw reads together with processed data were uploaded to NCBI with access No. GSE208657. The clean RNA-seq data were aligned against the TAIR10 reference genome using HiSAT2 (v.2.1.0, (61)) with following parameters, hisat2 -p 4 -x TAIR10indexed -1 sample_strand1.fq.gz -2 sample_strand2.fq.gz -S aligned_sample.sam. The aligned reads were assembled into transcripts according to the TAIR10 annotation (62, 63) using Stringtie (64). ∼ 99% overall alignment rate was reached for each sample. Differential expression analysis was done by comparing transcript levels between pairwise normalized samples using DEseq2 package in R (LFC ≥ ±0.58, padj ≤ 0.05) (65). Gene ontology was analyzed first against the whole *Arabidopsis* genome at TAIR Gene Ontology terms website, http://geneontology.org/ (66, 67), and which was then confirmed using total detected 22,622 (covers 88.7% of known genes in *Arabidopsis* genome) genes of the RNA-seq results on agriGO website http://systemsbiology.cau.edu.cn/agriGOv2/# (68). Figures were generated in R, with ComplexHeatmap package that was specifically used for Figure 3 B-D and 4 (69). Complete R scripts are available at SI Appendix, information S6.

### Motif discovery and enrichment analysis

The de novo motif discovery analysis was carried on the website (https://www.arabidopsis.org/tools/bulk/motiffinder/index.jsp). Frequencies of 6-mer motifs were compared between 1000 bp upstream sequences of each input gene and the current *Arabidopsis* genomic sequence set (33518 sequences). A total of 15 motifs containing the consensus SPL binding site, GTAC, were identified. A frequency-based sequence logo was generated through the weblogo, (https://weblogo.berkeley.edu/logo.cgi). To assess the enrichment of experimentally validated TF binding sites, we used the SEA (https://meme-suite.org/meme/tools/sea). 1000 bp upstream sequences of the 203 Adu/*r10* co-upregulated DEGs (Fig. 6 A) were extracted from TAIR. Shuffled input sequences were chosen as control sequences. The DAP motif database (47) were selected to identify the enriched motifs.

### qRT-PCR

Bacterial suspension with 1 × 10^8^ CFU/mL was infiltrated in *Arabidopsis* leaves. Leaf samples were harvested at 3 hours hpi. Each sample had 16-20 leaf discs derived from 6-10 individual plants. The leaf age of plants in soil was measured following the same standard as the above. For plants on ½ MS plates, 15-20 leaves were harvested per genotype per treatment as one biological rep. leaves 8-13 from 45-day-old of Col-0 with abaxial trichomes were used as adult leaves, leaves 1-2 from 18-day-old Col-0 were used as juvenile leaves. In comparison, leaves 1-2 from 21-day-old of *rSPL10* were used. For each bio-rep, three to six technical replicates were used in one qPCR run. RNA extraction was performed using Omega biotek EZNA plant RNA kit. The qPCR was performed in the Applied Biosystems QuantStudio 1 Real-Time PCR system with SYBR Green master mix (Applied Biosystems). PCR conditions were set as follows: 95°C for 5 mins, 40 cycles of 95°C for 15s, 56°C for 30s and 72°C for 20-30s. *SAND* (*AT2G28390*) or *TUB2* (*AT5G62690*) was used as reference genes. The relative expression was calculated using relative standard curve methods. Delta-delta CT was also used for calculation when knowing that the PCR efficiency for the primers was at least more than 95% in previous standard curve results. The oligonucleotides used here can be found in SI Appendix, Table 1.

### Estradiol-induced gene expression

The estradiol-inducible *MIM156* and *rSPL3* was constructed using a Gateway compatible version of the XVE system, as described by Brand et al., (70). The *MIM156* and *rSPL3* sequence were cloned into pMDC160. Transgenic plants were crossed to plants containing pMDC150-35S (70) and generated homozygous. The estradiol-inducible *rSPL10* was generated by cloning *rSPL10* into a modified pMDC7 vector tagged with Citrine and HA.

### HPLC-MS

Ten to sixty leaves 1-2 of Col-0 and *r10* each and three adult Col-0 leaves (plants sawn in soil) were collected as one biological repeat for each genotype or a developmental stage. 3 to 6 biological replicates from five (for adult tissues) to thirty (for juvenile tissues) individual plants were collected in total. The leaves were then lyophilized, powdered, and weighted for getting a comparable dry weight for all samples (weighting error ≤ 0.1 mg). Metabolites from 1.5 mg or 8.5 mg (in separate experimental repeats) of dry tissue of each biological replicate were extracted and were detected under Liquid chromatography-mass spectrometry (HPLC-MS). Amount of salicylic acid beta-glucoside (SAG) and salicylic acid (SA) were measured and calculated in the unit of nmole metabolite per gram of dry weight (nmole/g DW).

### ChIP-qPCR assay

The procedure and the preparation of reagents were modified from protocols (71-73). In brief, 1 gram of non-treated fully expanded adult leaf tissues were collected from the pro*SPL10::rSPL10-YFP* line. Leaves from 2-3 plants were harvested as one biological sample. Three biological replicates in total from two independent experiments were used. The tissue was crosslinked, and the chromatin was extracted and sonicated using nuclei extraction buffers and a bioruptor UCD-200 with chilling pump. 15 µL of post-sonicating chromatin solution was saved as the input. The remaining chromatin solution was immunoprecipitated by using GFP-trap Magnetic Agarose (ChromoTek, cat no. gtma). Diluted input and DNA eluted after IP were used for qPCR using primers at the indicated positions (Fig. 6 B; SI Appendix, Table 1). We used the following formula to calculate the estimated CT value for adjusting CT values of input and IP samples, ΔCt [normalized ChIP] = (Ct [Input] – Log2 (dilution factor for Input)) – ((Ct [ChIP] – Log2 (dilution factor for ChIP)), and the final output % input = 100 * 2 ^ (ΔCt [normalized ChIP] (73).

### GUS staining assay

Plants carrying pro*PR2*::GUS were harvested at six week after planting. The GUS solution was prepared as following and was vacuum infiltrated into plants, 0.1 M NaPO_4_, pH 7.0, 10 mM EDTA. 0.1% Triton X-100, 1 mM K_3_Fe(CN)_6_, 2 mM X-Gluc (X-Gluc was dissolved in N, N-DMF and made fresh). After 24 h incubation at 37°C, the staining solution was replaced with 70% ethanol. Tissues were washed several times with 70% ethanol until the chlorophyll in leaves was cleared. The GUS-stained leaves were imaged using a dissecting microscope (VWR).

## Supporting information

Rscripts

tableS1_seeds_primers

tableS2_replicates_in_Fig1257

tableS3_Adu_DEGs

tableS4_Juv_DEGs

tableS5_r10_DEGs

tableS6_GOs

## Acknowledgement

We thank Scott Poethig from University of Pennsylvania to provide *rSPL* and *spl* mutants. We thank Khadijeh Mozaffari, Scott Harding and Chung-Jui Tsai of the Plant Metabolomics Laboratory at the University of Georgia for analytical assistance. We thank Jovana Mijatovic for providing feedback on the manuscript. Y.L. conceptualized the project. Y.L. and H.L. designed experiments. H.L. performed experiments. P.A. and K.F. participated repeating experiments in Figure 2 A and Figure 6 B, respectively. H.L., Q.P. Y.Y and Y.L. analyzed data. H.L. and Y.L. wrote the manuscript. Funding for the UPLC-Q-TOF was provided by the United States Department of Agriculture, National Institute of Food and Agriculture, Equipment Grant Program award no. 2021-70410-35297. This project is supported by NIH R35GM143067 to Y.L.

---

**Figure S1.**
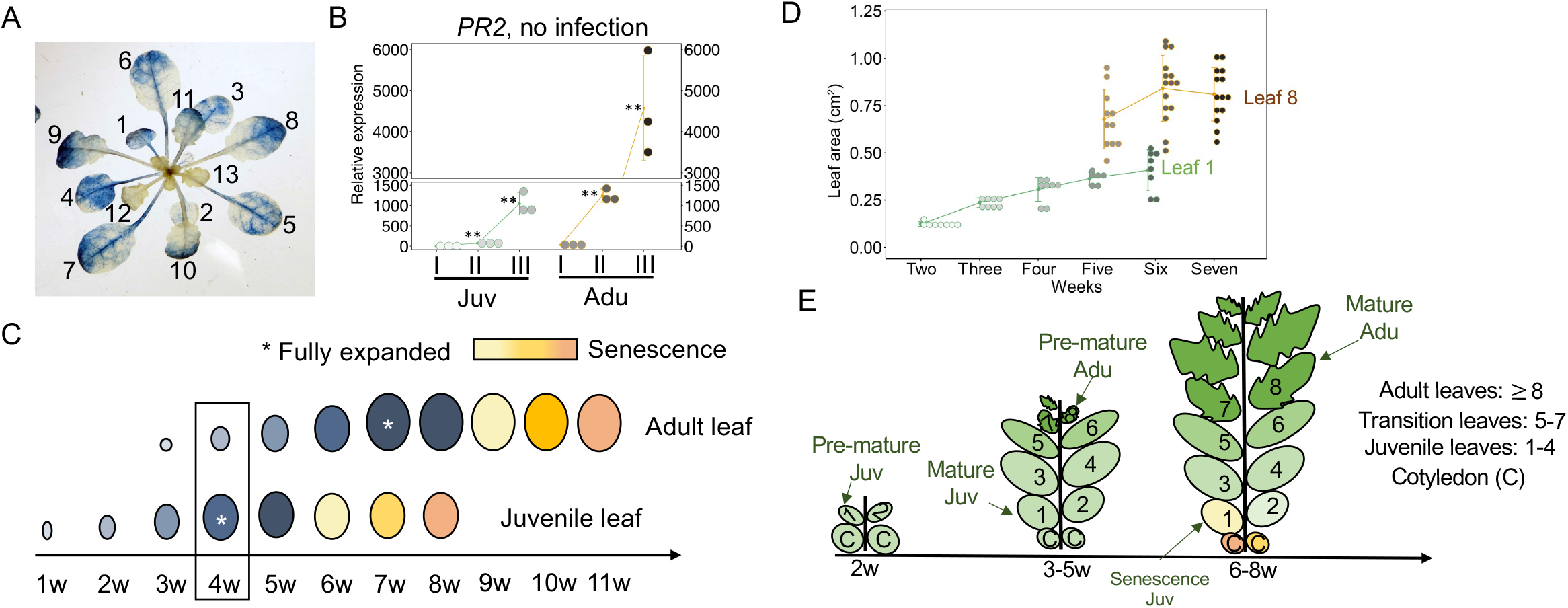
Ontogenic change of SA response and sampling approach for examining ARR_vpc_. **A**, ontogenic-associated promoter activities of pro*PR2*::GUS in an uninfected plant. Note the high activity in fully expanded juvenile leaves (1-7) and low in young adult leaves (12-13). The indicated leaf numbers were based on the order of the leaves on shoot. **B**, the incremental expression of *PR2* gene spanning the expansion of juvenile and adult leaves. I, premature leaves. II, intermediate premature leaves. III, mature leaves. **C**, An outline of the ontogenic maturation of a juvenile leaf and an adult leaf. The boxed region indicates distinct ontogenic age of juvenile and adult leaves from the same plant; asteroids indicate juvenile and adult leaves of the same ontogenic age. **D**, The quantified leaf expansion rate in juvenile and adult leaves. Leaf areas were quantified and normalized through Fiji software. **E**, a cartoon depiction of shoot development and leaf maturation that are concurrent during the vegetative phase change.

**Figure S2.**
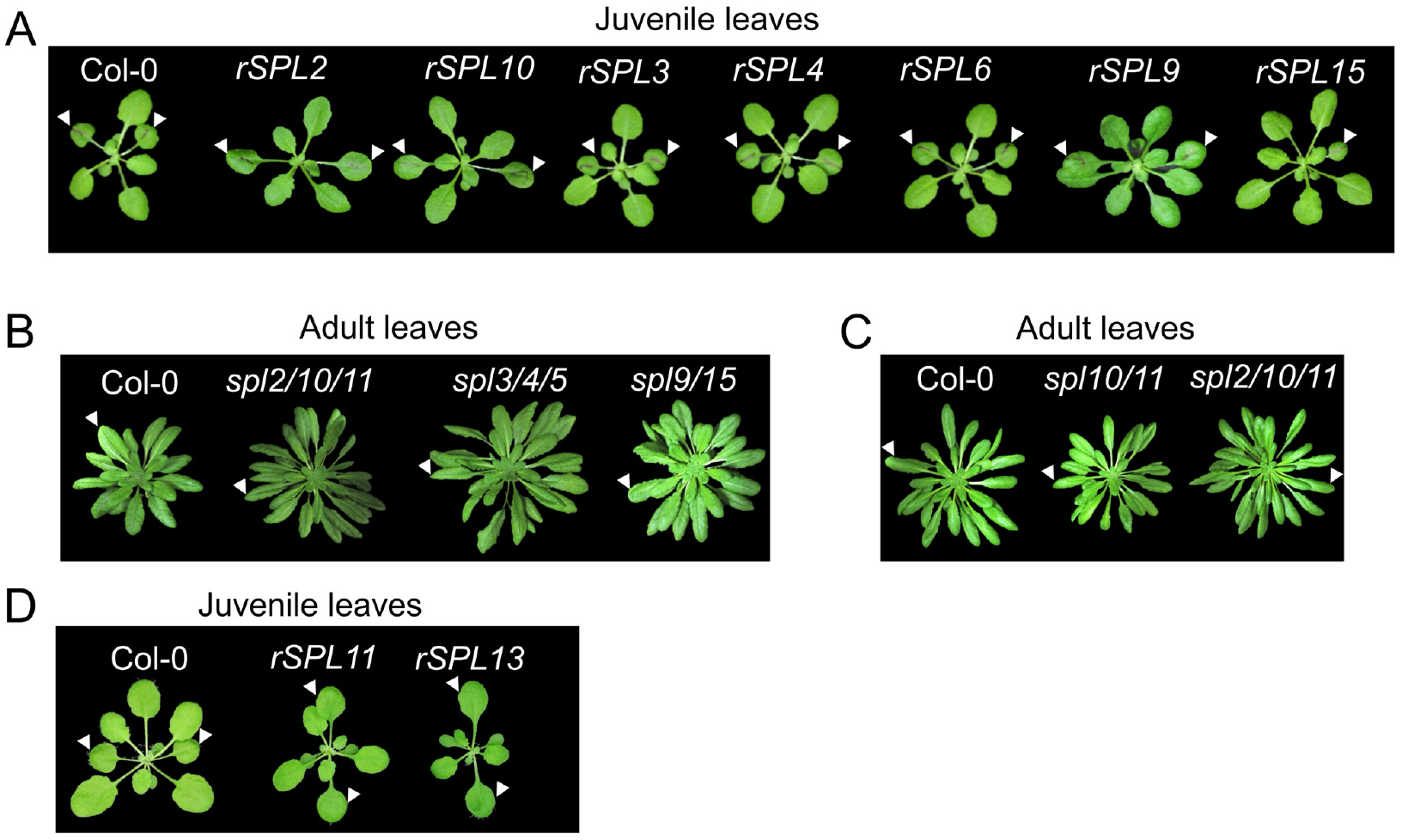
Developmental phenotypes of *rSPLs* and *spl* mutants. **A and D**, arrows indicate leaf 1-2 of juvenile Col-0 and *rSPLs*. **B-C**, arrows indicate representative adult leaves of Col-0 and *spl* mutants.

**Figure S3.**
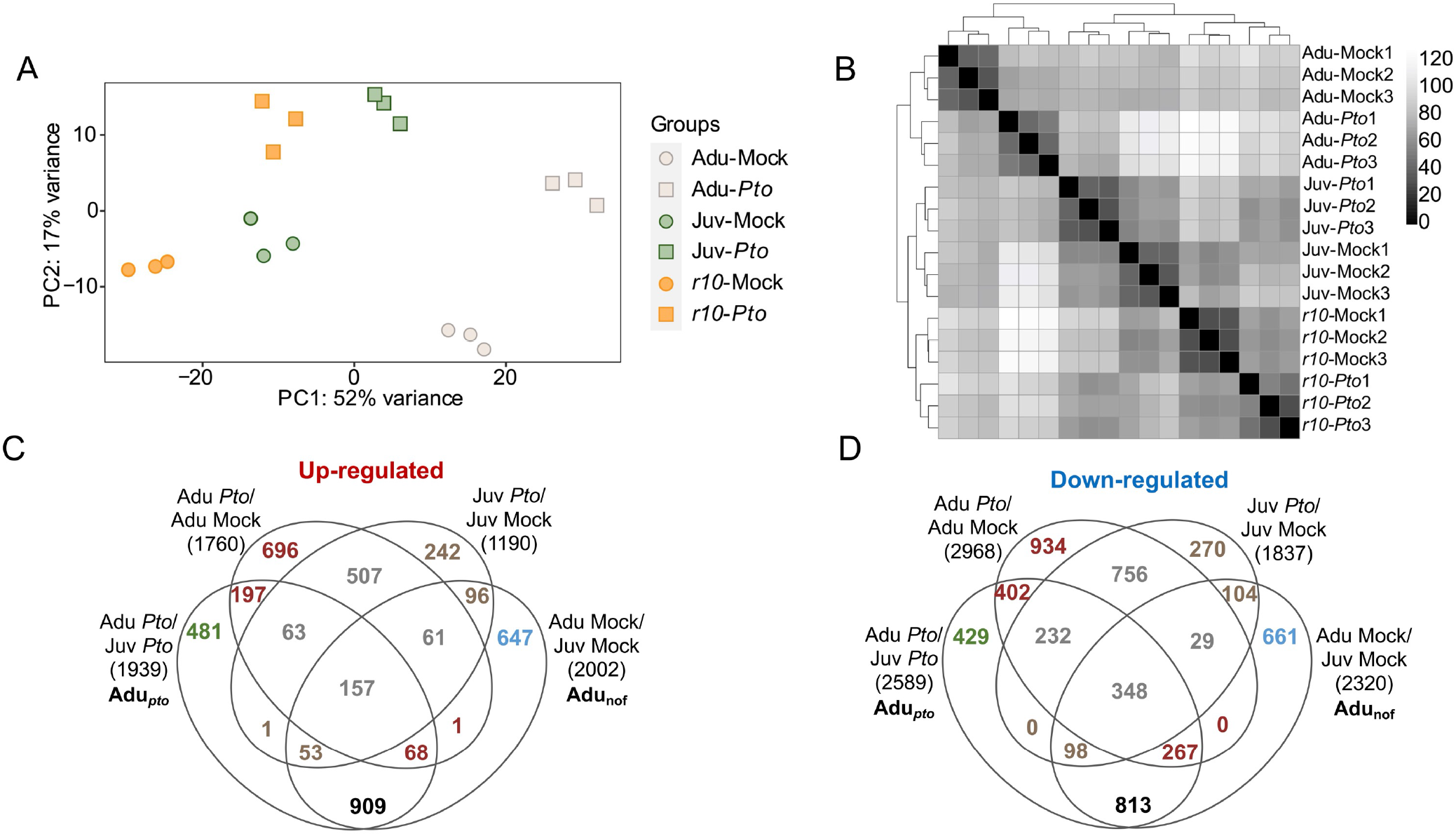
Quality control of RNA-seq datasets and Venn diagrams of adult *vs* juvenile DEGs. **A**, principal component analysis showed that the effect of age and genotype likely explained 52% variance of the samples, and *Pto* infection effect likely explained 17% variance of the samples. **B**, the sample-to-sample distance matrix showed that biological replicates (Mock1-3 and *Pto*1-3) were well correlated (in close distance) within per genotype per treatment. The column names of the matrix are in the same order as the row names--from the “Adu-Mock1” (the first on the left) to the *“r10-Pto3*” (the first on the right). **C**, venn diagrams of Adu-DEGs generated from the indicated pair-wise comparisons. Color-coding of numbers, adult-specifically *Pto*-triggered DEGs (red), Juvenile-specifically triggered (brown), commonly triggered in both adults and juveniles, i.e., shared (grey), and overlap DEGs between Adu_nof_ and Adu_*pto*_ (black). Green numbers indicate the 20.1% synergistic DEGs that mentioned in the main text. Blue numbers refer to DEGs that were Adu_nof_ but not Adu_*pto*_. Adult preferentially *Pto-*triggered DEGs are listed in SI Appendix Table S3.

**Figure S4.**
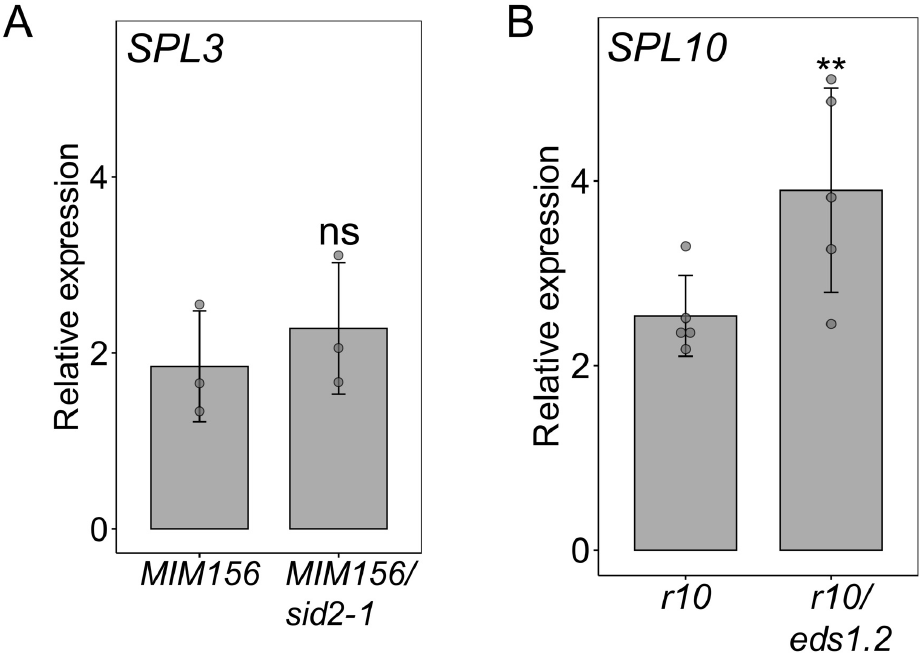
Expression of *SPL3* and *SPL10* in *MIM156/sid2-1* and *r10/eds1*.*2*. A, *SPL3* was used as an indicator of *MIM156* function. No difference was observed in *MIM156* and *MIM156/sid2-1*. B, *SPL10* was expressed at comparable level in *rSPL10* and *rSPL10/eds1*.*2*. Student t test, ns, not significant, *, p < 0.05, **, p < 0.01.

**Figure S5.**
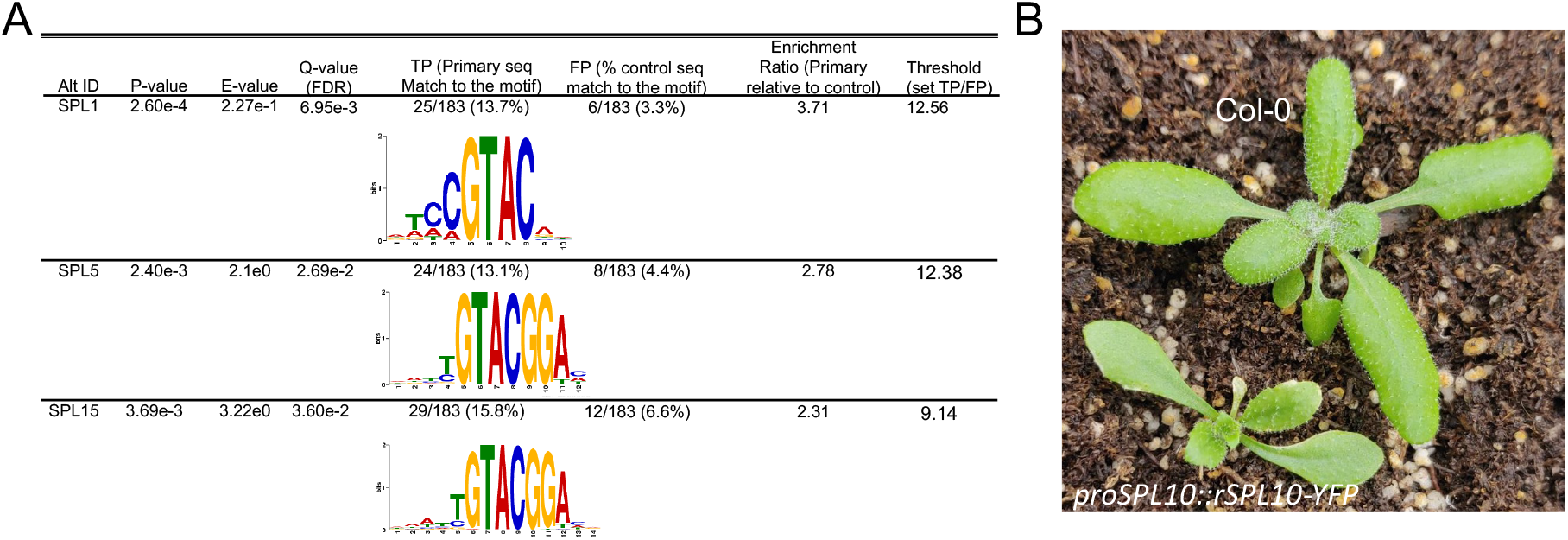
A table of enriched SPL-binding motifs within the upregulated 203 and plant phenotype of *proSPL10::rSPL10-YFP*. A. Adu/*r10*_nof_ co-regulon. Frequency-based DNA logos are shown for each enriched motif. The analysis was performed in the simple enrichment analysis-MEME Suite. Details were described in the method section. **B**. plant phenotype of proSPL10::rSPL10-YFP. Note the elongated leaves 1 and 2 in the transgenic plant.

**Figure S6.**
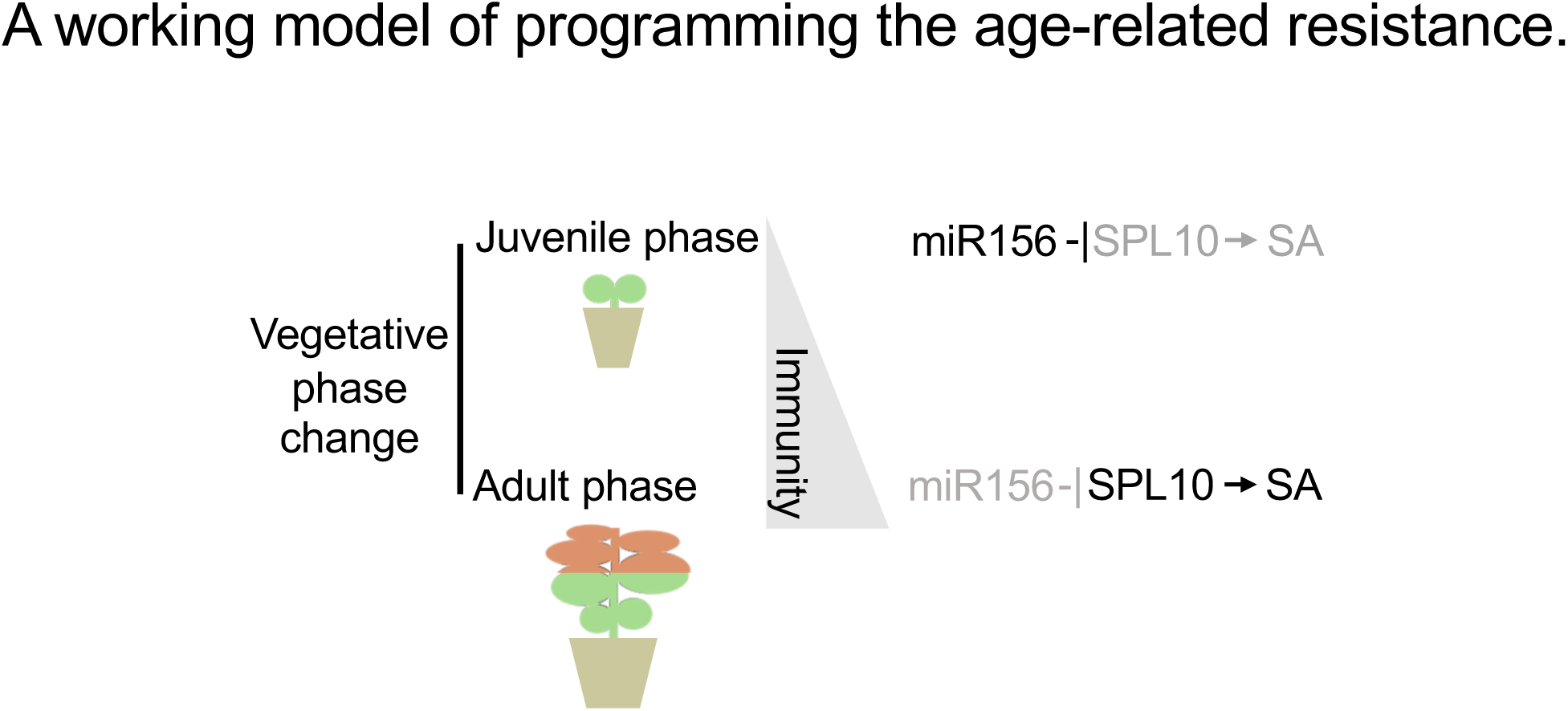
A model of miR156-SPL10 regulated age-related resistance during the vegetative phase change. In brief, miR156 suppressed the resistance in juvenile phase through inhibiting SPL10. The increased expression of *SPL10* followed by the decline of miR156 level gives rise to a high immune output in adult phase. That is achieved via promoting the expression of *PAD4* as well as enhancing expressions of components in SA biosynthesis and signaling pathways.

