## Supplementary material for "Distinct function of SPL genes in age-related resistance in *Arabidopsis*": Rscripts

### R scripts that are created and adapted for this paper.

If you adapt **following scripts** in your publications, we are appreciated if you can cite this paper and related references that are mentioned below or in the method section.

#### Figure 1B, 1D, 1F, 2B, 2D, and 7

**Dot-boxplots** used for displaying bacterial growth results.

```
library(ggplot2)
library(magrittr)
library(ggpubr)
library(RColorBrewer)
#Scripts used to generate each dot-boxplot are the same. Only Genotype and corresponding
#bacterial counts are vary between plots.
#read document
c = read.csv("/Users/IDxxxx/Desktop/figurex.csv")
head(c)
#Sort order of x axis
c$sample = factor(c$sample, levels=c("Genotype1", "Genotype2"))
#specify parameters for plot designs
theme_YL = theme(legend.position="top", legend.direction="horizontal",
panel.background = element_rect(fill = "white", colour = NA), panel.border = element_rect(fill = NA,
colour = "black"), panel.grid.major.y = element_line(colour = NA), panel.grid.major.x = element_line
(colour = NA), legend.text = element_text(size = 22.5), axis.text.y = element_text(size=22.5,
```

```
colour = "black"), axis.text.x = element_text(size=22.5, colour = "black"),
legend.title=element_text(size=22.5), axis.title.y=element_text(size=22.5),
axis.title.x=element_text(size=22.5))
#factorize the numeric variable "rep"
c$rep = factor(c$rep)
#set up the dot-boxplot
p= ggplot(c, aes(x=sample, y=bacterial_growth)) + geom_boxplot(aes(color=Days), fill=NA, alpha=1.0,
width=0.7, outlier.size=0, outlier.stroke=0, lwd=0.7, position=position_dodge(width=0)) +
geom_dotplot(aes(fill=rep, color=Days), alpha = 1.0, binaxis="y", binwidth = 0.15,
stackdir = "center", position=position_dodge(width=0.5)) + ylab("Log10"~CFU/cm^2) +
xlab("Genotype") + ylim(1, 8)
#call out the plot in R
p + scale_color_brewer(palette="Set1") + scale_fill_brewer(palette="Greys") + theme_YL +
theme(legend.position="top") + guides(fill = guide_legend(nrow = 1), color = guide_legend(nrow = 1))
#save the plot in .pdf format. Width may be vary depending on the number of genotypes in the plot.
ggsave("/Users/IDxxxx/Desktop/Col_vs_inMIM156_final.pdf", width= 5.00, height = 6.61)
```

### Figure 3B-C

**DE analysis** used for generating DE tables.

```
library(IRanges)
library(GenomicRanges)
library(GenomeInfoDb)
library(SummarizedExperiment)
library(MatrixGenerics)
library(matrixStats)
library(Biobase)
library(S4Vectors)
library(stats4)
```

```
library(BiocGenerics)
library(parallel)
library(DESeq2)
library(ggplot2)
library(coseq)
#Scripts used for each pair-wise comparison are the same as below, except certain samples were added
#or omitted in specific comparisons.
coldata <- read.csv("/Users/IDxxxx/Desktop/11pheno_master.csv", row.names=1)
coldata
Treatment
assembled_RLH10    r10_Juv_Mock
assembled_RLH11    r10_Juv_Mock
assembled_RLH13 r10_Juv_DC3000
assembled_RLH14 r10_Juv_DC3000
assembled_RLH16 r10_Juv_DC3000
assembled_RLH17    Col_Adu_Mock
assembled_RLH19    Col_Adu_Mock
assembled_RLH2     Col_Juv_Mock
assembled_RLH20    Col_Adu_Mock
assembled_RLH22 Col_Adu_DC3000
assembled_RLH23 Col_Adu_DC3000
assembled_RLH24 Col_Adu_DC3000
assembled_RLH3     Col_Juv_Mock
assembled_RLH4     Col_Juv_Mock
assembled_RLH6  Col_Juv_DC3000
assembled_RLH7  Col_Juv_DC3000
assembled_RLH8  Col_Juv_DC3000
assembled_RLH9     r10_Juv_Mock
countdata <- read.csv("/Users/IDxxxx/Desktop/10gene_count_matrix.csv", header=TRUE, row.names=1)
#The head of countdata is too large to display here. Ideally, the same arrangement shown in the
#.csv file should be seen in R.
head(countdata)
```

```
countdata <- as.matrix(countdata)
coldata$Treatment = factor(coldata$Treatment)
dds <- DESeqDataSetFromMatrix(countData = countdata, colData = coldata, design = ~Treatment)
keep <- rowSums(counts(dds)) >= 10
dds <- dds[keep,]
dds$Treatment <- relevel (dds$Treatment, ref = "Col_Adu_Mock")
dds <- DESeq(dds)
res <- results(dds)
write.csv(res,"/Users/IDxxxx/Desktop/merge_genelevel_DESeq2_21023_SMviiiCA.csv",row.names=TRUE)
baseMeanPerLvl <- sapply( levels(dds$Treatment), function(lvl) rowMeans( counts(dds,normalized=TRUE)
[,dds$Treatment == lvl] ))
res2 <- merge(baseMeanPerLvl, res, by=0, all=TRUE)
write.csv(res2,"/Users/IDxxxx/Desktop/merge_genelevel_DESeq2_afmeanmerged_21023_SMviiiCA.csv",
row.names=TRUE)
```

**The two heatmaps** used for displaying expression profile of Adu-DEGs. Note that the term "sdy" (steady state) shown in the following steps referred to the "nof" that is in the main text and figures.

```
#referencing source, https://jokergoo.github.io/ComplexHeatmap-reference/book/
library(grid)
library(ComplexHeatmap)
library(circlize)
#figure 3B
#read the .csv document with info for making the annotation bar
otter <- read.csv("/Users/lh10509/Desktop/fig3b1_sdy_reordered_cols38.csv")
head(otter)
Gene_ID        c1       c2     g1     g2   g3
1 AT4G23180 2.9200016 2.997575 Adusdy Adupto con
2 AT2G15090 2.6200517 1.942765 Adusdy Adupto con
3 AT1G64390 2.1095687 1.836339 Adusdy Adupto con
4 AT2G21660 1.3348047 1.723803 Adusdy Adupto con
```

```
5 AT5G64860 0.8680323 1.448265 Adusdy Adupto con
6 AT4G14400 2.5170900 1.386720 Adusdy Adupto con
#color-code the heatmap according to the LFC range, -9~9
col_fun = colorRamp2(c(-9, 0, 9), c("#377EB8", "#FFFFFF", "#E41A1C"))
#define the annotation bar. con refers to 909+813 DEGs that are not Pto-triggered
#but already differently expressed at steady state when comparing adult vs juv leaves.
row_ha = rowAnnotation(bar= otter$g3, col= list(bar= c("con" = "#000000", "#N/A" = "#FFFFFF")))
#read the .csv document that will be used for making the main heatmap body
mat <- read.csv("/Users/lh10509/Desktop/fig3b1_map_sdy_reordered_cols38.csv")
mat_matrix <- as.matrix(mat[, -c(1)])
rownames(mat_matrix) <- mat$Gene_ID
head(mat_matrix)
#c1=Adu-M/Juv-M and c2=Adu-P/Juv-P
c1        c2
AT4G23180 2.9200016 2.997575
AT2G15090 2.6200517 1.942765
AT1G64390 2.1095687 1.836339
AT2G21660 1.3348047 1.723803
AT5G64860 0.8680323 1.448265
AT4G14400 2.5170900 1.386720
#make the map, note that the column order is being sorted here.
p=Heatmap(mat_matrix, name = "LFC", col = col_fun, column_order = sort(colnames(mat_matrix)),
show_column_dend = FALSE, row_split = factor(otter$g3, levels= c("con", "#N/A")),
clustering_method_rows = "complete", clustering_distance_rows = "euclidean",
cluster_row_slices = FALSE, width = unit(3, "cm"), height = unit(8, "cm"),
heatmap_legend_param = list(direction = "horizontal"), right_annotation = row_ha)
#save the annotated heatmap in .pdf format
pdf(file="/Users/IDxxxx/Desktop/sdy_ARR_con.pdf")
draw(p, heatmap_legend_side = "top")
dev.off()

#figure 3C
```

```
#Same library packs were used.
#Attach and read the .csv document with info for making annotation bars
otter <- read.csv("/Users/IDxxxx/Desktop/inducible_ARR.csv")
head(otter)
Gene_ID          c3         c4      g1   g2
1 AT4G31200 -0.32935925 -0.5777023 Adu_spe #N/A
2 AT5G41610 -0.02427914 -0.5777470 Adu_spe #N/A
3 AT2G28830 -0.22187559 -0.5791842 Adu_spe #N/A
4 AT5G17710 -0.29095029 -0.5794031 Adu_spe #N/A
5 AT1G69730 -0.01182052 -0.5795746 Adu_spe #N/A
6 AT3G57080 -0.34364007 -0.5802866 Adu_spe #N/A
#color-code the heatmap according to the LFC range, -9~9
col_fun = colorRamp2(c(-9, 0, 9), c("#377EB8", "#FFFFFF", "#E41A1C"))
#read the .csv document that will be used for making the main heatmap body
mat <- read.csv("/Users/IDxxxx/Desktop/inducible_ARR_map.csv")
mat_matrix <- as.matrix(mat[, -c(1)])
rownames(mat_matrix) <- mat$Gene_ID
head(mat_matrix)
# c3=Juv-P/Juv-M, c4=Adu-P/Adu-M
c3          c4
AT4G31200 -0.32935925 -0.5777023
AT5G41610 -0.02427914 -0.5777470
AT2G28830 -0.22187559 -0.5791842
AT5G17710 -0.29095029 -0.5794031
AT1G69730 -0.01182052 -0.5795746
AT3G57080 -0.34364007 -0.5802866
#colorcoding annotation bars that will be on the right of the heatmap.
row_ha = rowAnnotation(bar= otter$g1, bar1= otter$g2, col= list(bar= c("Adu_spe" = "#960200",
"Adu_spe_pto" = "#960200", "Juv_spe_pto" = "#A88157", "pre" = "#004896",
"Common_pto" = "#969696", "Juv_spe" = "#A88157", "Common" = "#969696"),
bar1= c("Adupto" = "#000000", "#N/A" = "#FFFFFF")))
#make the map
```

```
p=Heatmap(mat_matrix, name = "LFC", col = col_fun, column_order = sort(colnames(mat_matrix)),
show_column_dend = FALSE, row_split = factor(otter$g1, levels= c("Adu_spe_pto", "Adu_spe",
"Juv_spe_pto", "Juv_spe", "pre", "Common_pto", "Common")), clustering_method_rows = "complete",
clustering_distance_rows = "euclidean", cluster_row_slices = FALSE, width = unit(3, "cm"),
height = unit(8, "cm"), heatmap_legend_param = list(direction = "horizontal"),
right_annotation = row_ha)
#save the map in .pdf format
pdf(file="/Users/IDxxxx/Desktop/inducible_ARR.pdf")
draw(p, heatmap_legend_side = "top")
dev.off()
```

### Figure 3D and 4C

**A heatmap variant** for displaying GO terms with multiple DEG categories.

```
library(grid)
library(ComplexHeatmap)
library(circlize)
#figure 3D and 4B used the same script except the input GO terms and fold enrichment values are
#different.
#read the .csv document that will be used to assign each GO term a name.
otter <- read.csv("/Users/IDxxxx/Desktop/combined_GOs_short.csv")
#define color codes for the heatmap
col_fun = colorRamp2(c(-9, 0, 9), c("#377EB8", "#FFFFFF", "#E41A1C"))
#read the .csv document for mapping out fold enrichment values with colors defined as above.
mat <- read.csv("/Users/IDxxxx/Desktop/combined_GOs_short_map.csv")
mat_matrix <- as.matrix(mat[, -c(1)])
rownames(mat_matrix) <- mat$Go_Terms
head(mat_matrix)
#The input GO terms and fold enrichment values here are for figure 3D. c1=Adusdy, c2=shared,
```

```
#c3=Adu-spe in Adupto.
c1    c2    c3
response to heat                                 0.00  4.33 4.76
Maturation of 5.8S-rRNA from tricistonic rRNA transcript 6.94  0.00 0.00
respiratory burst involved in defense response       0.00 21.46 0.00
defense response to bacterium                        2.19  4.23 3.03
defense response to fungus                           2.16  4.21 0.00
response to salicylic acid                           3.28  2.98 0.00
# set up the heatmap
p=Heatmap(mat_matrix, rect_gp = gpar(col = "black", lwd = 2), name = "Fold enrichment", col = col_fun,
column_order = sort(colnames(mat_matrix)), show_column_dend = FALSE, show_row_dend = FALSE,
clustering_method_rows = "complete", clustering_distance_rows = "euclidean",
cluster_row_slices = FALSE, width = unit(3, "cm"), height = unit(16, "cm"),
heatmap_legend_param = list(direction = "horizontal"), row_names_gp = gpar(fontsize = 20))
#save the map in .pdf format
pdf(file="/Users/IDxxxx/Desktop/figure_3D.pdf")
draw(p, heatmap_legend_side = "top")
dev.off()
```

### Figure 4A and 4B

**Heatmaps** used for displaying co-regulated Adu/*r10* DEGs.

```
library(grid)
library(ComplexHeatmap)
library(circlize)
#Scripts for 4A and 4C are the same except the input files are different.
#read the .csv document
otter <- read.csv("/Users/IDxxxx/Desktop/coadur10_sdy_full_ep.csv")
head(otter)
```

```
    Gene_ID      Adu      r10 Label
1 AT4G21650 7.012627 8.366337  #N/A
2 AT5G15160 7.672442 7.579840  #N/A
3 AT4G31840 5.951223 6.904511  #N/A
4 AT3G23890 6.033988 6.888020  #N/A
5 AT2G40610 6.549540 6.294097  #N/A
6 AT3G60470 6.622070 6.194341  #N/A
#define color codes for the heatmap
col_fun = colorRamp2(c(-9, 0, 9), c("#377EB8", "#FFFFFF", "#E41A1C"))
#read the .csv document for mapping out fold enrichment values with colors defined as above.
mat <- read.csv("/Users/IDxxxx/Desktop/coadur10_sdy_full_ep_map.csv")
mat_matrix <- as.matrix(mat[, -c(1)])
rownames(mat_matrix) <- mat$Gene_ID
               Adu      r10
AT4G21650 7.012627 8.366337
AT5G15160 7.672442 7.579840
AT4G31840 5.951223 6.904511
AT3G23890 6.033988 6.888020
AT2G40610 6.549540 6.294097
AT3G60470 6.622070 6.194341
head(mat_matrix)
#define annotations, EDS1 and PAD4.
row_ha = rowAnnotation(bar= otter$Label, col= list(bar= c("EDS1" = "#960200", "PAD4" = "#000000",
"#N/A" = "#FFFFFF")))
#use PAM clustering and manually define the cluster numbers and orders.
pa = cluster::pam(otter_matrix, k = 4)
#set up the map.
p=Heatmap(otter_matrix, name = "LFC", column_order = sort(colnames(otter_matrix)), col = col_fun,
show_column_dend = FALSE, row_split = factor(paste0("reg", pa$clustering),
levels= c("reg4", "reg2", "reg3", "reg1")), cluster_row_slices = FALSE, width = unit(3, "cm"),
height = unit(12, "cm"), heatmap_legend_param = list(direction = "horizontal"),
right_annotation = row_ha)
```

```
#save the map in .pdf format
pdf(file="/Users/IDxxxx/Desktop/figure_4A.pdf")
draw(p, heatmap_legend_side = "top")
dev.off()
```

#### Figure 5B

**Volcano plots** for displaying different expression profile of upregulated SA markers or randomly detected genes between comparisons of Adu *vs* Juv or *r10 vs* Juv, under steady state or *Pto* infection state.

```
library(ggplot2)
library(ggrepel)
# read the .csv input file
c <- read.csv("/Users/IDxxxx/Desktop/Adusdy_SAupm.csv")
head(c)
    Gene_ID      LFC     padj DE     shape
1 AT3G60470 6.622070 6.43e-05 Up AT3G60470
2 AT2G14610 6.297867 1.98e-09 Up     other
3 AT4G10500 6.174617 4.60e-28 Up     other
4 AT1G75040 5.637495 7.56e-13 Up       PR5
5 AT3G57240 5.630796 5.91e-76 Up       BG3
6 AT5G55450 5.118302 3.36e-13 Up  ATLTP4.4
#sort orders for the DE column.
c$DE = factor(c$DE, levels=c("Up", "No", "Down"))
theme_YL = theme(legend.position="top", legend.direction="horizontal", panel.background
= element_rect(fill = "white", colour = NA), panel.border = element_rect(fill = NA,
colour = "black"), panel.grid.major.y = element_line(colour = NA),
panel.grid.major.x = element_line(colour = NA), legend.text = element_text(size = 22.5),
axis.text.y = element_text(size=22.5, colour = "black"), axis.text.x = element_text(size=22.5,
colour = "black"), legend.title=element_text(size=22.5),
```

```
axis.title.y=element_text(size=22.5),
axis.title.x=element_text(size=22.5))
#make the plot. After making this plot. I made a seperate plot without "aes(shape=shape), size=4.5"
#and then manually annotated the four SA markers in the plots with positions of different shapes
#from previous plots as references. Alternatively, can use enhanced volcano plot to combine
#annotations within plots,
#https://bioconductor.org/packages/devel/bioc/vignettes/EnhancedVolcano/inst/doc/EnhancedVolcano.html.
ggplot(data=c, aes(x=LFC, y=-log10(padj), col=DE)) +
        geom_point(aes(shape=shape), size=4.5) +
        theme_YL +
        scale_color_manual(values=c("#E41A1C", "#969696", "#377EB8")) +
        geom_vline(xintercept=c(-0.58, 0.58), col="#E41A1C") +
        geom_hline(yintercept=-log10(0.05), col="#E41A1C") +
        ylim(-1, 120) + xlim(-8, 10)
ggsave("/Users/IDxxxx/Desktop/Adusdy_SAupm_shape.pdf", width= 4.50, height = 4.50)
```

**Hypergeometric test** for statistics of the overlaps between core up- or down-regulated SA markers with Adu and r10 upregulated DEGs.

```
#Hygrogemetric test for figure 5 B
#Use Adu and r10 upregulated DEGs under steady state or Pto infection state to test against
#the SA core markers
> library(GeneOverlap)
> Adur10_DEGs_up <- read.csv("/Users/lh10509/Desktop/Adur10_up.csv")
> head(Adur10_DEGs_up)
Adu_sdyDEGs_up Adu_PtotriDEGs_up r10_sdyDEGs_up r10_PtotriDEGs_up
1       AT5G35425          AT5G62165      AT4G21650          AT5G59310
2       AT5G62165          AT1G28290      AT2G35637          AT2G37362
3       AT1G31710          AT5G35920      AT1G44830          AT2G13660
4       AT4G24540          AT4G24540      AT5G15160          AT5G37300
5       AT3G23290          AT4G18870      AT4G13420          AT2G19200
6       AT1G33840          AT5G26660      AT2G30933          AT3G26200
```

```
> SA_cores <- read.csv("/Users/lh10509/Desktop/SA_cores.csv")
> head(SA_cores)
core_SAmarkers_up core_SAmarkers_down
1         AT1G01340           AT1G01320
2         AT1G01560           AT1G02800
3         AT1G02230           AT1G03270
4         AT1G02360           AT1G03310
5         AT1G02450           AT1G07440
6         AT1G02850           AT1G08550
> gom.obj <- newGOM(Adur10_DEGs_up, SA_cores, 27000)
> drawHeatmap(gom.obj)
> getMatrix(gom.obj, name="pval")
                      core_SAmarkers_up core_SAmarkers_down
Adu_sdyDEGs_up         9.565852e-77        7.962293e-01
Adu_PtotriDEGs_up      2.442415e-42        9.992267e-01
r10_sdyDEGs_up         4.563681e-86        2.580411e-07
r10_PtotriDEGs_up      6.740559e-38        3.945944e-02
> getMatrix(gom.obj, "odds.ratio")
                      core_SAmarkers_up core_SAmarkers_down
Adu_sdyDEGs_up             6.222783           0.8342993
Adu_PtotriDEGs_up          4.271533           0.3904240
r10_sdyDEGs_up             7.005481           2.5856009
r10_PtotriDEGs_up          4.298375           1.5189362
#Use random set of genes derived from each pair-wise comparison as negative controls to test against
#the SA core markers
> random_inAdur10 <- read.csv("/Users/lh10509/Desktop/random_inAdur10.csv")
> gom.obj <- newGOM(random_inAdur10, SA_cores, 27000)
> drawHeatmap(gom.obj)
> getMatrix(gom.obj, name="pval")
                      core_SAmarkers_up core_SAmarkers_down
Random_AduMvsJuvM        0.33145524          0.01609447
Random_AduPvsJuvP        0.03405089          0.16849757
```

```
Random_r10MvsJuvM        0.44506624          0.08518882
Random_r10PvsJuvP        0.15635748          0.00608961
> getMatrix(gom.obj, "odds.ratio")
                      core_SAmarkers_up core_SAmarkers_down
Random_AduMvsJuvM              1.168751            2.289072
Random_AduPvsJuvP              1.649798            1.573256
Random_r10MvsJuvM              1.074692            1.808924
Random_r10PvsJuvP              1.358965            2.533501
```

### Figure 5C-E and 6B-C

**Bar-dotplots** for displaying endogenous accumulations of SA and SAG as well as results from (ChIP-) qPCR assays.

```
#figure 5C
library(dplyr)
library(ggplot2)
library(magrittr)
library(ggpubr)
c = read.csv("/Users/IDxxxx/Desktop/fig5b.csv")
head(c)
  Genotype      conc reps cd
1      Juv  4.185519    1 SA
2      Juv  4.585482    1 SA
3      Juv  7.560252    1 SA
4      r10  7.951860    1 SA
5      r10 11.215278    1 SA
6      Adu 11.023260    1 SA
c$cd = factor(c$cd, levels=c("SAG", "SA"))
c$reps = factor(c$reps)
#specify parameters for plot designs
```

```
theme_YL = theme(legend.position="top", legend.direction="horizontal",
panel.background = element_rect(fill = "white", colour = NA), panel.border = element_rect(fill = NA,
colour = "black"), panel.grid.major.y = element_line(colour = NA), panel.grid.major.x = element_line
(colour = NA), legend.text = element_text(size = 22.5), axis.text.y = element_text(size=22.5,
colour = "black"), axis.text.x = element_text(size=22.5, colour = "black"),
legend.title=element_text(size=22.5), axis.title.y=element_text(size=22.5),
axis.title.x=element_text(size=22.5))
#generate the plot
p= ggbarplot(c, x = "Genotype", y = "conc", add = c("mean_sd"), fill= "cd", alpha = 0.60,
position = position_dodge(width=0.8), ylab="nmole/g DW") + ylim (-0.5, 30) +
geom_dotplot(aes(x=Genotype, y=conc, fill=cd, color=reps), alpha = 0.70, binaxis="y",
binwidth = 0.8, stackdir = "center", position=position_dodge(width=0.8))
p + theme_YL + theme(legend.position="top") + scale_fill_manual(values =
c("#000000", "#737373"), 2) + scale_color_manual(values = c("black")) + guides(fill =
guide_legend(nrow = 1), color = guide_legend(nrow = 1))
ggsave("/Users/IDxxxx/Desktop/fig5d.pdf", width= 5.69, height = 9.61)

#figure 5D and 6B-C. Similar settings were used with different values of the relative expression
# (or % input in the case of figure 6B) and with different color codes.
#Same library packs as the above were used.
c = read.csv("/Users/IDxxxx/Desktop/fig5d.csv")
head(c)
      Genotype relative_expression     Genes reps
1      Adu_Col               2.397 AT3G60470    1
2      Adu_Col               1.494 AT3G60470    1
3      Adu_Col               1.788 AT3G60470    1
4 Adu_spl21011               0.923 AT3G60470    1
5 Adu_spl21011               1.083 AT3G60470    1
6      Adu_Col               1.959       BG3    1
c$reps = factor(c$reps)
c$Genotype = factor(c$Genotype, levels=c("Adu_Col", "Adu_spl21011"))
c$Genes = factor(c$Genes, levels=c("AT3G60470", "BG3", "ATLTP4.4", "PR5"))
```

```r
p= ggbarplot(c, x = "Genotype", y = "relative_expression", add = c("mean_sd"), color="Genes",
fill="Genes", alpha=0.60, palette = get_palette(c("#000000", "#737373", "#710C0D", "#1E4564")
, 4), position = position_dodge(width=0.8), ylab="Relative expression") +
geom_dotplot(aes(x=Genotype, y=relative_expression, fill=Genes, color=Genes), alpha = 0.70,
binaxis="y", binwidth = 0.2, stackdir = "center", position=position_dodge(width=0.8)) + ylim(-1, 8)
#theme_YL is the same as the above
p + theme_YL + theme(legend.position="top") + guides(fill = guide_legend(nrow = 2),
color = guide_legend(nrow = 2))
ggsave("/Users/IDxxxx/Desktop/fig5c.pdf", width= 6.69, height = 6.61)

#figure 5E
#add two more libraries in addition to those above
library(patchwork)
library(ggbreak)
c = read.csv("/Users/IDxxxx/Desktop/fig5d.csv")
head(c)
  Genotype relative_expression           Genes reps
1      Juv               0.181 putative_DUF247    1
2      Juv               5.652 putative_DUF247    1
3      r10              65.765 putative_DUF247    1
4      r10             110.957 putative_DUF247    1
5      r10              79.547 putative_DUF247    1
6      Adu               9.103 putative_DUF247    1
c$reps = factor(c$reps)
c$Genotype = factor(c$Genotype, levels=c("Juv", "r10", "Adu"))
#AT3G60470 = putative_DUF247
c$Genes = factor(c$Genes, levels=c("putative_DUF247", "BG3", "ATLTP4.4", "PR5"))
p= ggbarplot(c, x = "Genotype", y = "relative_expression", add = c("mean_sd"), color="Genes",
fill="Genes", alpha=0.60, palette = get_palette(c("#000000", "#737373", "#710C0D", "#1E4564")
, 4), position = position_dodge(width=0.8), ylab="Relative expression") + ylim (-1, 210) +
geom_dotplot(aes(x=Genotype, y=relative_expression, fill=Genes, color=Genes), alpha = 0.70,
binaxis="y", binwidth = 2, stackdir = "center", position=position_dodge(width=0.8)) +
```

```
scale_y_break(c(30, 60), scales = 0.8) + scale_y_break(c(90, 120), scales=1) +
scale_y_break(c(150, 180), scales=0.8)
#theme_YL is the same as the above
p + theme_YL + theme(legend.position="top") + guides(fill = guide_legend(nrow = 1),
color = guide_legend(nrow = 2))
ggsave("/Users/IDxxxx/Desktop/fig5e.pdf", width= 6.69, height = 6.61)
```
